# Convergent Agonist and Heat Activation of Nociceptor TRPM3

**DOI:** 10.1101/2025.01.23.634542

**Authors:** Sushant Kumar, Fei Jin, Sung Jin Park, Wooyoung Choi, Sarah I. Keuning, Richard P. Massimino, Simon Vu, Wei Lü, Juan Du

## Abstract

Detecting noxious heat is vital for survival, triggering pain responses that protect against harm^1,2^. The TRPM3 channel is a key nociceptor for sensing noxious heat and a promising therapeutic target for pain treatment and neurological disorders such as epilepsy^3–11^. Here, we functionally and structurally characterized TRPM3 in response to diverse stimuli: the synthetic superagonist CIM0216 ^Ref12^, the anticonvulsant antagonist primidone^13,14^, and heat^1,10,15^. Our findings reveal that TRPM3 is intrinsically dynamic, with its intracellular domain (ICD) sampling both resting and activated states, though strongly favoring the resting state without stimulation. CIM0216 binds to the S1–S4 domain, inducing conformational changes in the ICD and shifting the equilibrium toward activation. Remarkably, heat induces similar ICD rearrangements, revealing a converged activation mechanism driven by chemical compounds and temperature. This mechanism is supported by functional data showing that mutations facilitating the ICD movement markedly increase the sensitivity of TRPM3 to both chemical and thermal signals. These findings establish a critical role of the ICD in temperature sensing in TRPM3, a mechanism likely conserved across the TRPM family. Finally, we show that primidone binds to the same site as CIM0216 but acts as an antagonist. This study provides a framework for understanding the thermal sensing mechanisms of temperature-sensitive ion channels and offers a structural foundation for developing TRPM3-target therapeutics for pain and neurological disorders.

## Main

Temperature profoundly influences the activity of numerous proteins in the human body, shaping our ability to respond to a wide spectrum of thermal stimuli, including noxious cold, cool, warm and noxious heat^1,7,10,16–29^. Detecting extreme temperature such as noxious heat is a critical survival mechanism that is closely linked to pain signaling, as heat-induced responses trigger protective mechanisms to prevent tissue damage^1,2^. Pain perception begins at the sensory neuron level, where specialized ion channels detect harmful stimuli and transduce signals to the nervous system^2,18,30,31^.

Among these channels, TRPM3, a member of the transient receptor potential (TRP) family, functions as a thermosensor activated by noxious heat^1,3,10,15,32,33^. It is a voltage-dependent, Ca^2+^-permeable, non-selective cation channel expressed in sensory neurons, including the dorsal root and trigeminal ganglia. TRPM3 shows substantial activation at approximately 35 °C, with markedly increased activity between 40 °C and 45 °C, within the range of noxious heat^10^. Beyond temperature sensing, TRPM3 also acts as a chemosensor and its activity is modulated by various ligands, including neuroactive steroid pregnenolone sulfate (PregS)^8,34^, synthetic superagonist CIM0216^Ref^^12^, and antagonist primidone^13,14^, which enhance or diminish TRPM3 activity, consequently altering pain effect.

Deletion of TRPM3 in mice causes clear deficits in their avoidance responses to noxious heat and impairs the development of inflammatory heat hyperalgesia, highlighting the channel’s relevance to pain signaling^10^. Indeed, TRPM3 has been implicated in various pain-related conditions, including inflammatory thermal hyperalgesia and female migraine^11,35–37^, making it a promising but underexplored target for pain management. Moreover, gain-of-function mutations in TRPM3 are associated with epilepsy, and its inhibitor primidone—a first-generation barbiturate type antiepileptic drug—mitigates seizures^4,6,13,14,38–41^. However, beyond primidone, selective drugs targeting TRPM3 for therapeutic applications are lacking. Despite its essential roles in pain signaling and neurological disorders, the mechanisms underlying TRPM3’s temperature- and ligand-induced activation and inhibition remain poorly understood. Addressing these knowledge gaps could pave the way for the development of targeted therapies for TRPM3 - related conditions.

To explore how TRPM3 responds to diverse stimuli, we determined its structures under apo condition and in response to heat, as well as ligands including CIM0216, and primidone. Structural dynamics analysis provided insights into how these stimuli activate TRPM3, and revealed the molecular determinants of temperature sensing and ligand binding. Furthermore, the findings highlight a converged mechanism through which heat and chemical ligands modulate the channel via shared structural transitions.

### Functional Characterization of rabbit TRPM3

To investigate the polymodal gating of TRPM3, particularly its activation by heat, we identified rabbit TRPM3 (rTRPM3) as an ideal candidate due to its robust biochemical stability following thermal exposure—a common destabilizing factor for purified proteins. The rTRPM3 shares 99.3% sequence identity with the human TRPM3 splice variant, which lacks the C-terminal domain (residues 1351–1732 in human TRPM3) of unknown function located immediately after the pole helix. Electrophysiological experiments confirmed the thermo- and chemosensitivity of rTRPM3. Heating or exposure to the endogenous agonist PregS and the superagonist CIM0216 induced reversible activation of an outwardly rectifying current (Extended Data Fig. 1a–c). The half-maximal effective concentrations (EC_50_) of CIM0216 and PregS were determined to be 0.7 µM and 22 µM at 80 mV membrane potential, respectively (Extended Data Figs. 1d, 2a), aligning with prior findings^8,12^. Furthermore, the antagonist primidone suppressed rTRPM3 currents induced by PregS. The estimated half-maximal inhibitory concentration (IC_50_) for primidone was approximately 5 µM at 80 mV membrane potential (Extended Data Fig. 2j), consistent with earlier studies on mouse TRPM3^Ref13^.

### TRPM3 activation by superagonist CIM0216

Unlike chemical ligands that can be directly observed, temperature is intangible, making it difficult to study temperature-dependent TRPM3 activation. Therefore, we first used CIM0216— a superagonist of TRPM3 that elicits robust, double-rectifying currents^12^ (Extended Data Fig. 1b)—to study the structural and dynamic changes during channel activation, aiming to establish a foundation for addressing the challenges of studying temperature-dependent activation.

We determined the cryo-EM structure of rTRPM3 in the presence of CIM0216 at an overall resolution of 3.0 Å (Extended data Figs. 3–5). The overall assembly closely resembles other TRPM channels^42^ (Fig. 1a, left panel). Each protomer consists of a transmembrane domain (TMD) and an intracellular domain (ICD), which includes the N-terminal melastatin homology regions (MHR1–4) and the C-terminal rib and pole helices. The TMD includes the S1–S4 domain and the pore domain (S5–S6), followed by the amphiphilic TRP helix. Notably, a small turret between the pore loop and S6 protrudes into the extracellular space. A prominent T-shaped density, hewing the shape of a CIM0216 molecule, was observed within the S1–S4 domain (Fig. 1a, right panel), interacting with residues on the transmembrane helices S1, S2, and S3, as well as the TRP helix, through both hydrophobic and hydrophilic contacts (Fig. 1b). Alanine substitutions at these key residues diminished or abolished CIM0216-evoked currents (Fig. 1c), confirming the functional relevance of these interactions. The horizontal part of the T-shaped CIM0216 fits into the binding pocket of the S1–S4 domain, while the tip of the vertical part penetrates deeper into its core, filling the inner cavity and forcing a 90-degree flip of the side chain of Y888 away from the domain center (Fig. 1b).

**Figure 1:**
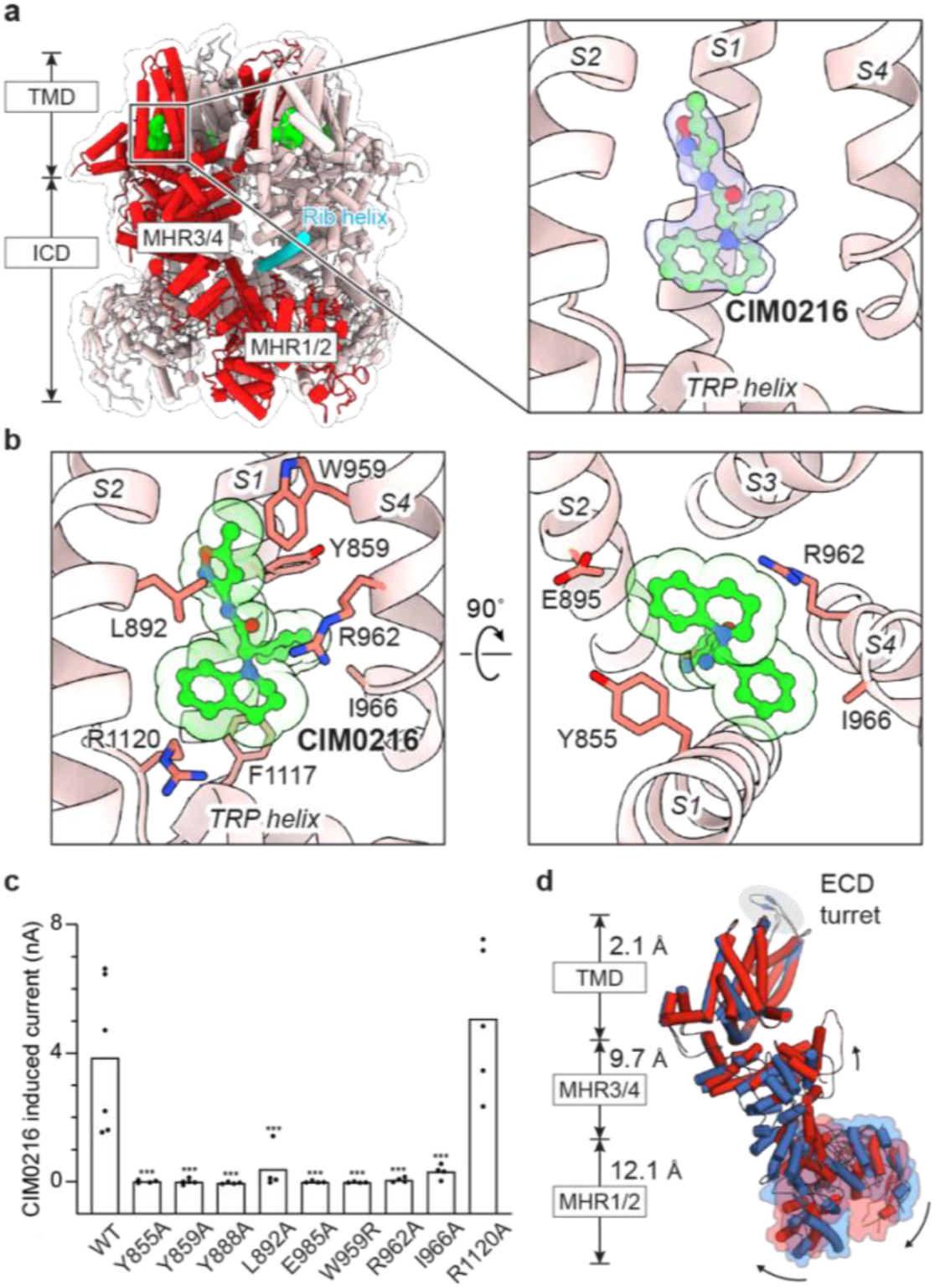
TRPM3 activation by superagonist CIM0216. **a**, Structure of TRPM3 in complex with CIM0216, shown in cartoon representation and viewed parallel to the membrane. CIM0216 (green) is fitted into the cryo-EM density map (right panel). One subunit is highlighted in red, and the rib helix from the adjacent subunit is highlighted in cyan. **b**, Close-up view of the CIM0216 binding site, detailing key interactions. **c**, Whole-cell currents measured in tsA cells overexpressing wild-type TRPM3 or CIM0216 binding site mutants at 22 °C. A voltage protocol was applied every 5 s to monitor current changes: −140 mV for 50 ms, ramped to +140 mV over 200 ms, and held at +140 mV for 50 ms. CIM0216-induced current amplitudes at +140 mV were calculated by subtracting the steady-state current recorded before CIM0216 application from the steady-state current recorded after CIM0216 application. Each point represents a single cell (from left to right, n = 6, 4, 5, 4, 4, 4, 4, 4, 4, 5), with bars indicating the mean. Statistical analysis was performed using one-way ANOVA with Bonferroni’s post hoc test (***P < 0.001). **d**, Structural comparison of CIM0216-bound activated state (red) and apo resting state (blue), superimposed using the TMD (residues 848–1119). Root-mean-square deviations (r.m.s.d.) are shown for individual domains. Black arrows indicate the approximate direction of domain movements.

**Figure 2:**
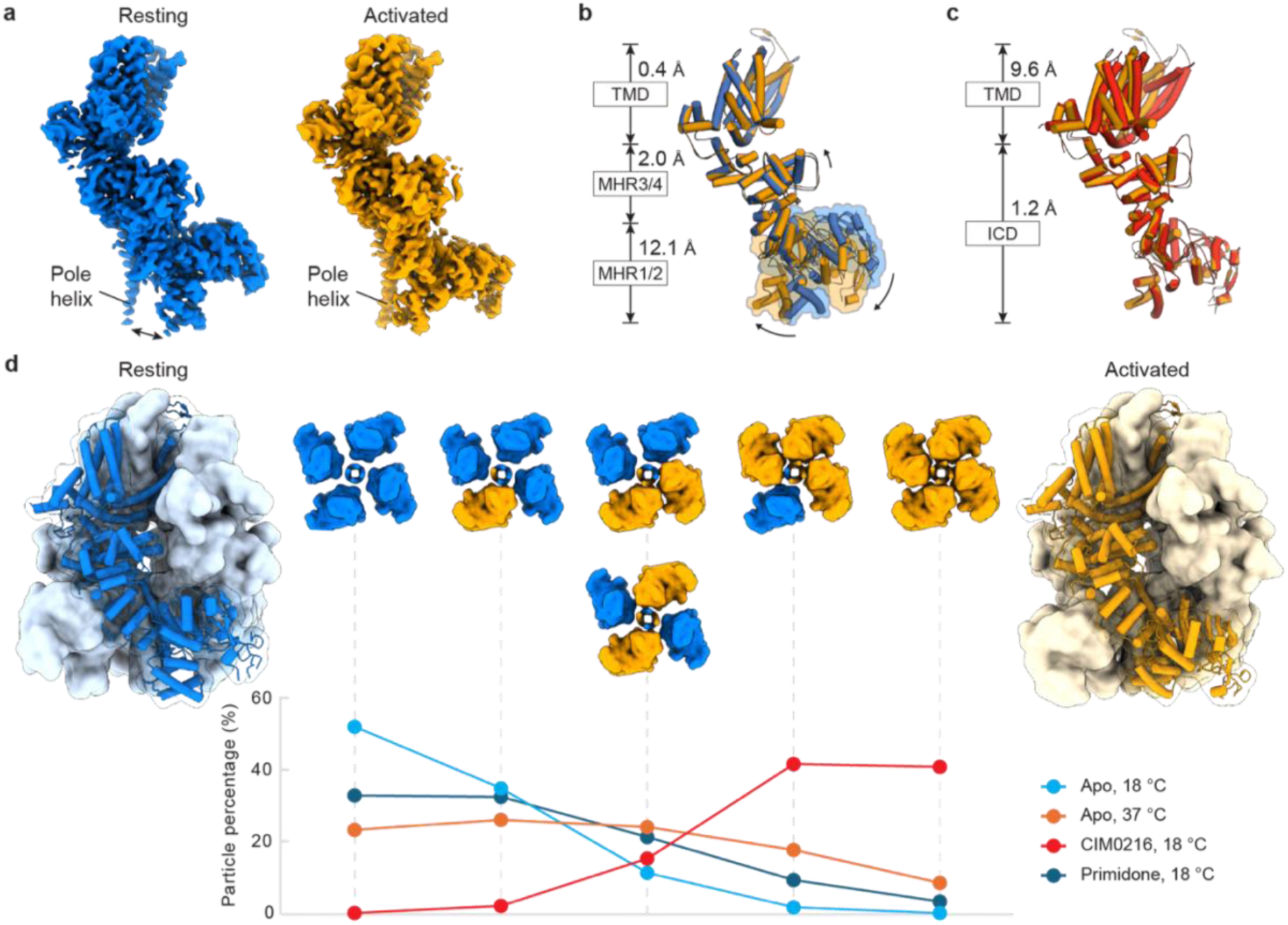
TRPM3 conformational dynamics. **a**, Cryo-EM density map of the apo resting state and apo activated state (right). The black double arrow in the left panel indicates the separation between the MHR1/2 domain and the C-terminal pole helix in the resting state, which come closer in the activated state. **b**, Structural comparison of apo resting state (blue) and apo activated state (orange), superimposed using the TMD. R.m.s.d. are shown for individual domains. Black arrows indicate the approximate direction of domain movements. **c**, Structural comparison of apo activated state (orange) and CIM0216-bound activated state (red), superimposed using the ICD (residues 1–747). R.m.s.d. are shown for individual domains. **d**, Structures of homotetrameric TRPM3 composed of four resting state subunits (left, blue) and four activated state subunits (right, orange). These structures are shown in surface representation, with one subunit displayed in cartoon and viewed parallel to the membrane. The six possible tetrameric compositions, shown in the middle panel, illustrate the coexistence of resting and activated subunits under different conditions (ligand and temperature), along with their respective occupancies. The percentages of the two compositions with two resting and two activated subunits (orthogonal and parallel positions) are combined. These compositions are displayed in surface representation viewed from the extracellular side, with resting subunits in blue and activated subunits in orange.

**Figure 3:**
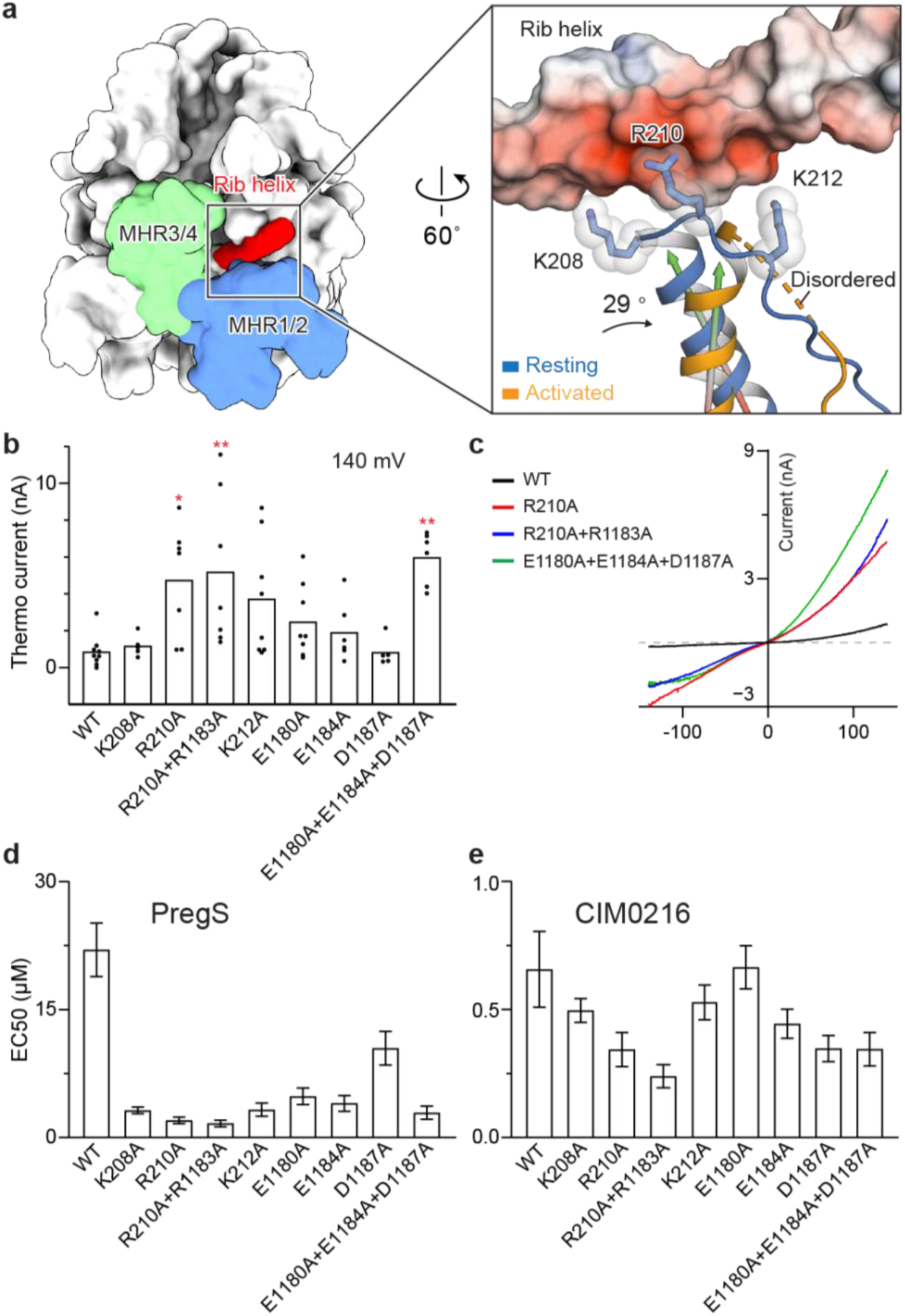
The MHR1/2 domain and rib helix interaction as a shared determinant of agonist- and heat-induced activation. **a**, Left: structure of TRPM3 in white surface representation, with the MHR1/2 and MHR3/4 domains highlighted in blue and green, respectively, and the rib helix of the adjacent subunit highlighted in red. Right: close-up comparison of the interaction between the positively charged loop in the MHR1/2 domain and the rib helix in the resting and activated states. The structures of the two states are superimposed using the rib helix (residues 1163 – 1192). The rib helix is shown as a surface representation colored according to the electrostatic surface potential (from –5 to +5 kT e^−1^, red to blue). The loop and its preceding helix in the MHR1/2 domain are shown in cartoon representation for the resting state (blue) and activated state (orange). The three positively charged residues are displayed in stick representation with transparent spheres. The angle between the helices in the resting and activated subunits is indicated. **b**, Whole-cell currents measured in tsA cells overexpressing wild-type TRPM3 or mutants at the interface between the MHR1/2 domain and the rib helix. A voltage protocol was applied every 5 s to monitor current changes: −140 mV for 50 ms, ramped to +140 mV over 200 ms, and held at +100 mV for 50 ms. Currents were measured as the temperature increased from 22 °C to 37 °C. Thermo currents at 140 mV were calculated by subtracting the steady-state current at 22 °C from the steady-state current at 37 °C. Mean raw data are presented in Extended Data Fig. 9. Each point represents a single cell (from left to right, n = 9, 5, 7, 7, 8, 8, 6, 5, 6), with bars indicating the mean. Statistical analysis was performed using one-way ANOVA with Bonferroni’s post hoc test (*P < 0.05, **P < 0.01). **c**, I/V curves corresponding to panel (b) for wild-type TRPM3 and three mutants, measured after the currents reached a steady state. **d**, EC_50_ values for PregS in wild-type TRPM3 and variants from panel (b). Data are presented as mean ± s.e.m. (from left to right, n = 6, 5, 5, 5, 5, 5, 5, 5, 5). **e**, EC_50_ values for CIM0216 in wild-type TRPM3 and variants from panel (b). Data are presented as mean ± s.e.m. (from left to right, n = 8, 4, 4, 5, 5, 4, 4, 5, 4).

**Figure 4:**
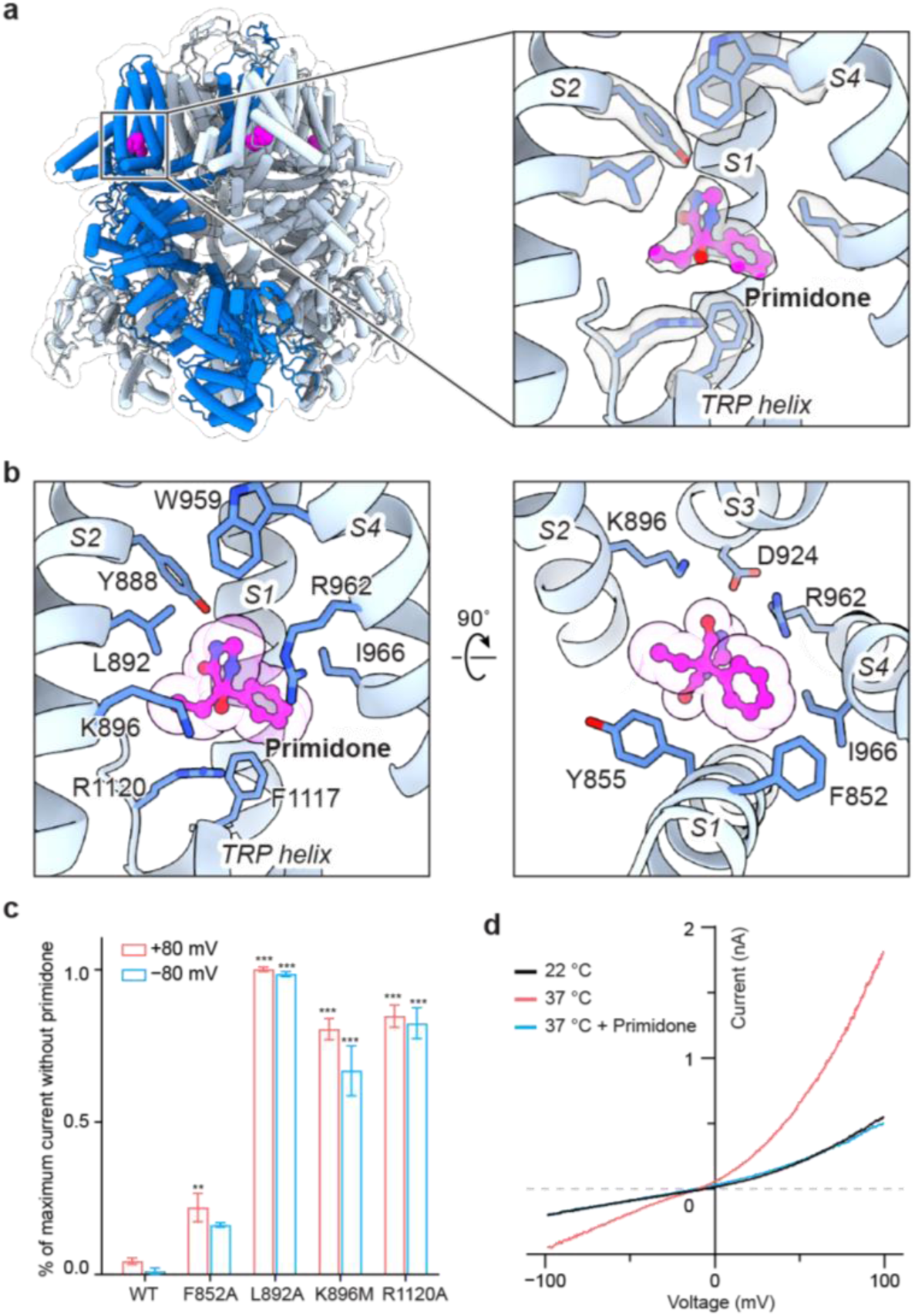
Primidone inhibition. **a**, Structure of TRPM3 in complex with primidone, shown in cartoon representation and viewed parallel to the membrane. Primidone (magenta) is fitted into the cryo-EM density map (right panel). One subunit is highlighted in blue. **b**, Close-up view of the primidone binding site, detailing key interactions. **c**, Ratio of whole-cell PregS-induced currents measured in the presence and absence of primidone for tsA cells overexpressing wild-type TRPM3 or primidone binding site mutants at 22 °C. A protocol was applied every 5 s to monitor current changes: 0 mV for 50 ms, switch to +80 mV for 50 ms, followed −80 mV for 50 ms, and a return to 0 mV for 50 ms. The ratio of remaining currents after the application of primidone was calculated using steady-state currents at +80 mV and −80 mV. Data are presented as mean ± s.e.m. (from left to right, n = 5, 3, 3, 3, 4). Statistical analysis was performed using one-way ANOVA with Bonferroni’s post hoc test (*P < 0.05, **P < 0.01, ***P < 0.001). **d**, I/V curves showing primidone inhibition of heat-activated whole-cell currents for wild-type TRPM3. The traces represent average currents at 22 °C, 37 °C, and 37 °C with primidone (n = 5, paired).

**Figure 5:**
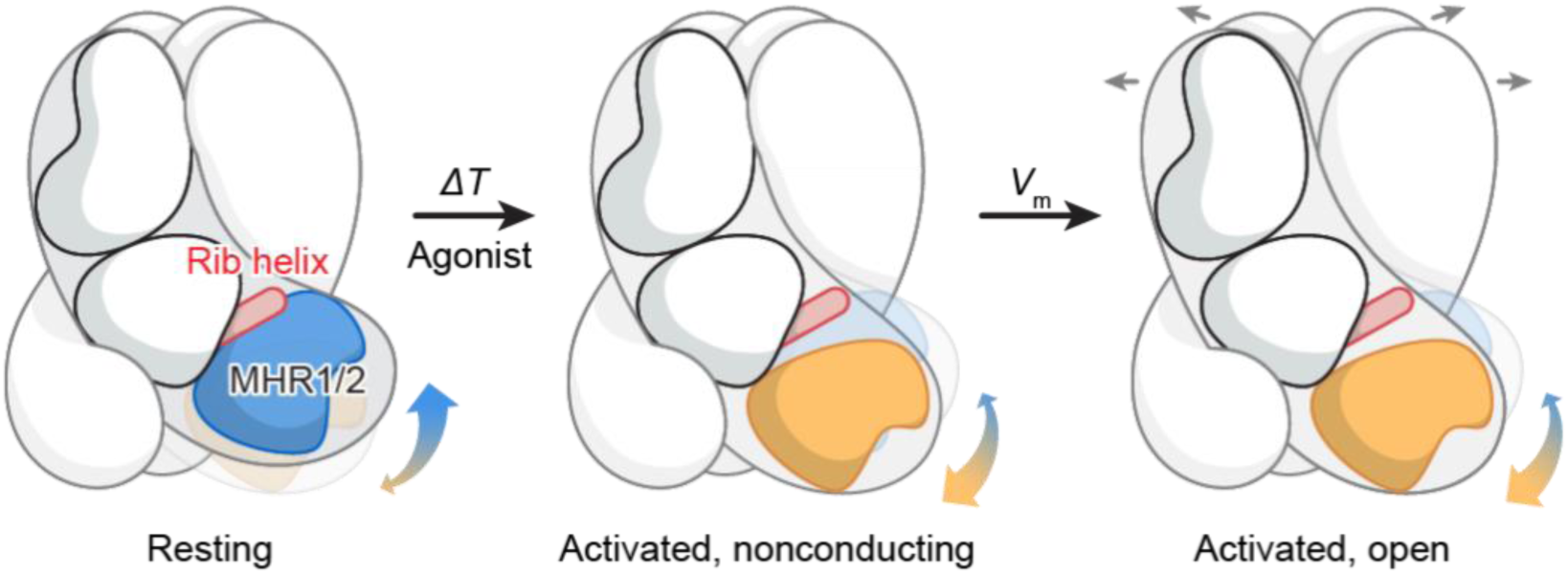
Converged activation mechanism by heat and agonist. Schematic representation of the polymodal activation of TRPM3 by chemical agonists and heat. Both stimuli induce a similar large-scale conformational rearrangement in the ICD, characterized by the disruption of the interface between the MHR1/2 domain and the adjacent rib helix. These changes may facilitate structural transitions in the TMD to promote the opening of the ion-conducting pore in response to membrane depolarization or other cellular cues. The structure of an activated, fully open state of TRPM3 remains to be determined.

The pore radius at the gate of the central conducting pore increases relative to the apo resting state of mouse TRPM3^Ref43^, however, it remains only slightly larger than a fully dehydrated sodium ion. This suggests that additional factors, such as membrane potential, are required to fully open the channel, consistent with the rectifying current induced by CIM0216 (Extended data Fig. 1b)^8,12^.

### Conformational dynamics of TRPM3 with CIM0216

Although the cryo-EM map of the TMD was of high quality, the ICD—particularly the distal MHR1/2—exhibited conformational heterogeneity. This observation aligns with previous findings on other TRPM channels, where the ICDs adopt distinct conformations that play a critical role in regulating properties such as voltage dependence and sensitivity to temperature and ligands^44,45^. Motivated by these insights, we analyzed the structure at the level of individual subunits, identifying two distinct subunit conformations: a major conformation representing 80% of subunits, and a minor conformation accounting for 20% (Fig. 1d; Extended data Fig. 3, text highlighted in red). Both conformations were bound to CIM0216. The minor conformation closely resembled the apo resting state of mouse TRPM3, with differences largely localized to the S1–S4 domain due to CIM0216 binding. In contrast, the major conformation displayed substantial changes, with a backbone root mean square deviation (r.m.s.d.) of ∼7 Å compared to the apo resting state of mouse TRPM3. The difference is even more pronounced when the TMD of a subunit is superimposed, as the MHR1/2 domain between the two states shows a remarkable backbone r.m.s.d. of 12 Å (Fig. 1d). These extensive structural rearrangements, particularly within the ICD, resemble the activation transition observed in TRPM4 upon heat and Ca ^2+^ binding^45^, suggesting that the major conformation represents a novel activated state induced by CIM0216.

In this activated state, CIM0216 binding elevates the TRP helix toward the extracellular side and causes its rotational movement along the pore axis, along with a shift in the S4 helix and S4–S5 linker. These regions, which are key gating elements in the TRP superfamily, mediate communication between the TMD and ICD, and within the TMD (linking the S1–S4 domain to the S5–S6 pore domain). Consequently, their movements propagate structural changes to both the ICD and the pore domain, driving extensive rearrangements throughout the protein (Fig. 1d).

To determine whether these conformational states exist exclusively in separate tetramers or co-exist within the same tetramer, we analyzed the composition of all tetrameric particles. Our analysis revealed six distinct tetrameric conformations with varying combinations of resting and activated subunits (Extended Data Fig. 3; Fig. 2d). Tetramers with either four or three activated subunits were predominant, while those with two activated subunits were less frequent. Tetramers containing one or no activated subunit were rare. These findings suggest that, even bound to a superagonist, TRPM3 channels exist in a dynamic equilibrium between resting and activated conformations.

### Heat shifts TRPM3 conformational dynamics

Building on these findings with CIM0216-induced activation, we next sought to investigate how heat activates TRPM3 in the absence of an agonist. To this end, we determined the cryo-EM structures of rTRPM3 at 18 °C and 37 °C, respectively (Extended data Figs. 4–7). We chose 37 °C for heat-induced study for two key reasons: functional data indicate that TRPM3 begins to activate at around 35 °C and shows substantial, but not maximal, activity at 37° C, resulting in a mixture of resting and activated conformations ideal for capturing temperature-dependent dynamics^10^. Additionally, temperatures exceeding 37 °C compromised the quality of cryo-EM data.

Despite achieving overall resolutions as high as 2.6 Å, we observed profound conformational heterogeneity in the ICDs, particularly the distal MHR1/2 domain, as indicated by the less well defined cryo-EM map in these regions. Structural analysis at the subunit level also revealed two distinct conformations (Fig. 2a; Extended data Fig. 6): a resting conformation, and a second conformation similar to the activated state observed with CIM0216, though differing in its TMD due to the absence of ligand binding (Fig. 2b, c). This second conformation likely represents an activated state in the absence of ligand binding. These findings align with the conformational selection model, which posits that proteins intrinsically sample a range of conformations— including both resting and activated states—and that external stimuli, such as temperature or ligands, shift the equilibrium toward specific conformations. Consistent with this model, our 37 °C data demonstrated that heat markedly altered the conformational distribution: at 18 °C, the resting conformation dominated (84%), whereas at 37 °C, the proportion of the activated conformation increased to 40% (Extended data Fig. 7, text highlighted in red).

At the tetrameric level, we also observed that resting and activated subunits co-exist within individual tetramers, forming six distinct combinations of resting and activated subunits (Fig. 2d). At 18 °C, tetramers with predominantly resting subunits (three or four) were most common, while tetramers with three or four activated subunits were rare. This distribution, combined with the lack of basal activity in TRPM3 at low temperatures in the absence of an agonist, suggests that channel activation likely requires at least three, if not all four, subunits to adopt the activated conformation. At 37 °C, the proportion of tetramers with three or four activated subunits increased 14-fold relative to 18 °C, rising from 2% to 27%. However, this is still markedly lower than the 83% observed with CIM0216-bound TRPM3 (Fig. 2d). This difference aligns with functional data showing that maximal heat-induced TRPM3 activity occurs at temperatures higher than 37 °C and that heat-induced TRPM3 currents at 37 °C are substantially smaller than those evoked by CIM0216^Ref10^ (Extended data Fig. 3a, b). Additionally, it should be noted that the temperature at which the cryo-EM sample was “frozen” was likely somewhat below the intended 37 °C due to instrument limitations, which could contribute to the lower observed distribution of activated states. Notably, the ion-conducting pore in tetramers with predominantly activated subunits remained closed, consistent with the fact that heat-induced TRPM3 opening is voltage-dependent and produces outwardly rectifying currents.

### Converged activation mechanism by heat and agonist

Structural comparisons between the resting and activated conformations—derived from all datasets mentioned above—revealed a consistent trend: the ICD undergoes a large counterclockwise inward rotation during the transition from resting to activated states when viewed from the intracellular side (Figs. 1d, 2b; Extended data Fig. 8a). A key and conserved change was observed at the interface between the MHR1/2 domain and the rib helix: these regions interact in the resting conformation but dissociate in the activated state (Fig. 3a; Extended data Fig. 8b), regardless of whether activation is driven by an agonist or heat.

The rib helix, which penetrates the ICD and forms multiple interactions with it, connects to the TRP helix and plays a critical role in transducing signals from the ICD to the TMD. In the resting conformation, the rib helix, containing 11 negatively charged residues, establishes electrostatic interactions with a positively charged loop in the MHR1 domain (Fig. 3a). In the activated conformation, however, this loop flipped away and became disordered, disrupting its interactions with the rib helix and coinciding with a rearrangement of the ICD. We hypothesized that this dissociation is linked to the conformational dynamics underlying heat- and agonist-dependent activation. To test this idea, we neutralized the charged residues involved in the interaction to potentially weaken it, reasoning that a weakened interaction would facilitate the structural transition from the resting to the activated state, thereby enhancing channel sensitivity to both temperature and agonists.

Consistent with this hypothesis, mutants designed to weaken these interactions not only produced markedly larger heat-induced currents but also altered the voltage sensitivity in electrophysiological experiments (Fig. 3b, c; Extended data Fig. 9). Notably, some mutants showed substantial inward currents at negative potentials, where the wild-type channel showed no conductance (Fig. 3c). Additionally, these mutants demonstrated dramatically enhanced activation by agonists, with up to 13-fold higher sensitivity to the endogenous agonist PregS and up to 3-fold higher sensitivity to the CIM0216, a highly potent superagonist that strongly favors the activated conformation, in dose-response experiments (Fig. 2d, Fig. 3d, e; Extended data Figs. 1d–l, 2a–i).

Together, our structural and functional analysis reveals that TRPM3’s inherent structural dynamics enable it to naturally sample both resting and activated states, providing flexibility for diverse activation stimuli. External stimuli, such as heat or agonists, bias this equilibrium toward the activated state, promoting channel activation. Both heat- and agonist-induced activation share a common structural transition in the ICD, specifically the dissociation of the MHR1/2 domain from the rib helix. This unified mechanism illustrates how distinct stimuli converge to activate TRPM3, with the ICD playing a central role in mediating channel activation.

### Primidone-induced inhibition

The anticonvulsant primidone suppresses pain and treats seizures by inhibiting TRPM3^13,14^. To investigate the binding site and mechanism of action of the primidone, we determined the structure of rTRPM3 in complex with primidone at a resolution of 2.8 Å (Extended data Figs. 5, 10). A prominent density was observed within the cavity of the S1–S4 domain, which closely matches the shape of the primidone molecule, allowing for an unambiguous fit (Fig. 4a). The binding was further confirmed by mutagenesis studies, where alanine substitutions at key residues markedly reduced or abolished the inhibitory effect (Fig. 4b, c).

Interestingly, despite primidone acting as an inhibitor and CIM0216 as an agonist, both compounds occupy the same cavity within the S1–S4 domain (Fig. 4a; Fig. 1a). A comparison of the primidone-bound and the CIM0216-bound structures reveals how their opposing effects arise. CIM0216, being a bulky molecule that inserts deeper into the S1–S4 domain, induces substantial movement in the TRP helices, which is sandwiched between the S1–S4 domain and the ICD. This movement propagates to the ICD, shifting the conformational equilibrium toward the activated state (Figs. 1d, 2d). Moreover, CIM0216 displaces the S4 helix and S4–S5 linker, thereby altering the conformation the S5–S6 pore domain. Together, these changes in the ICD and the pore domain may work allosterically to facilitate channel opening in the presence of membrane potential.

Primidone, by contrast, is smaller and fits snugly into the S1–S4 cavity without causing major conformational changes. Its binding primarily affects the TRP helix and S3 helix, along with localized adjustments in the side chains of residues involved in primidone binding. The small movement in the S3 helix, located at the periphery of the TMD facing the lipid bilayer, is unlikely to impact the overall protein conformation. By contrast, the shift in the TRP helix resembles that observed with CIM0216, albeit smaller, and is therefore expected to propagate to the ICD to a lesser extent. Consistent with this, structural analysis at the subunit level revealed a moderate increase in the population of activated subunits upon primidone binding, from 16% to 30% (Extended data Figs. 6, 10), accompanied by a proportional rise in tetramers containing three or four activated subunits (Fig. 2d).

Importantly, however, primidone binding does not alter the conformation of the S4 helix or S4– S5 linker, leaving the S5–S6 pore domain in the resting state. This effectively locks the channel in a non-conductive state, regardless of the conformation of the ICD. This inhibition mechanism aligns with electrophysiology data showing that primidone effectively suppresses heat-induced channel activation (Fig. 4d), which primarily drives conformational changes in the ICD. Additionally, primidone likely acts as a competitive antagonist when inhibiting CIM0216 - induced currents, as both compounds compete for the same binding site. Our findings demonstrate that two ligands share the same binding site yet have opposing effects because they exert distinct structural impacts, driving diverse mechanisms of TRPM3 modulation.

## Discussion

Effective pain management involves either reducing neuronal excitation or enhancing inhibition within the nervous system. Opioids, for instance, reduce neurotransmitter release presynaptically and hyperpolarize neurons postsynaptically^46^. However, their high risk of addiction highlights the need for alternative approaches. Targeting nociceptors, specialized sensory neurons that initiate pain signals, offers a promising strategy by modulating ion channels like TRPM3, which detect noxious stimuli^47^. This study provides a comprehensive analysis of the polymodal gating mechanisms of TRPM3, revealing its ability to respond to diverse stimuli, including heat and chemical ligands, through distinct and shared structural transitions. We identified key molecular determinants for the binding of superagonist CIM0216 and antagonist primidone. Interestingly, CIM0216 and primidone, despite their opposing effects, share the same binding site within the S1–S4 domain. Structural comparisons revealed that their distinct effects arise from differences in size and shape, dictating their ability to allosterically drive conformational changes in the intracellular and pore domains. This contrast underscores the versatility of the S1 –S4 domain as a binding site capable of accommodating diverse ligands while producing distinct functional outcomes.

We demonstrated that heat and the agonist CIM0216 promote TRPM3 activation by shifting the conformational equilibrium from the resting state toward the activated state (Fig. 5). Despite their distinct modes of action—CIM0216 exerting its effect through binding to the TMD, while heat directly influences the ICD—both stimuli converge on a shared structural transition. This transition involves the dissociation of the MHR1/2 domain from the rib helix of the adjacent subunit within the ICD, highlighting a converged mechanism by which distinct stimuli activate the channel.

Building on our findings, we explored TRPM3’s thermal sensing properties in comparison to other temperature-sensitive TRP channels such as TRPM4 and TRPV^45,48,49^. In TRPM4, channel activation at high temperature involves significant intersubunit rearrangements in the ICD^45^. These large-scale structural reorganizations require substantial energy, making heat alone insufficient to drive activation. Instead, calcium, an additional activation factor, is required to facilitate the structural rearrangement of the ICD, and works synergistically with heat to activate the channel. In TRPM3, heat-induced conformational changes in the ICD primarily involve the dissociation of the MHR1/2 domain from the rib helix. Notably, the electrostatic interaction between these regions appears relatively limited, relying mainly on a short positively charged loop and the rib helix (Fig. 3a). This limited interaction likely provides sufficient flexibility for the ICD to undergo thermal activation without requiring additional energy input from another factor like calcium. This distinction may reflect the role of heat as a direct activation stimulus in TRPM3, compared to its role as a modulator cooperating with calcium in TRPM4. Importantly, our studies on TRPM4 and TRPM3 highlight the critical role of the ICD as a key sensor of temperature changes, likely representing a conserved mechanism within the TRPM family. In contrast, TRPV channels employ distinct structural mechanisms for thermal activation. In TRPV1, heat induces global yet subtle conformational changes, encompassing the ICD and the TMD ^48^. In TRPV3, thermal activation involves structural changes in the S2–S3 linker within the TMD, as well as rearrangements in both the N- and C-termini^49^. These differences underscore the diversity of thermal sensing mechanisms within and across protein families, likely tailored to their distinct physiological roles.

A key unresolved question is the binding site of the endogenous agonist PregS and its mechanism of action. Despite extensive efforts, we could not confidently identify the PregS binding site, likely due to its structural similarity to cholesterol or cholesterol hemisuccinate (CHS), a known TRPM3 modulator^34^. CHS was indispensable for isolating TRPM3 in our system, complicating the structural observation of PregS binding as CHS and PregS may compete for the same site(s). Nevertheless, our structures provided clues about potential PregS binding regions. Within the TMD, six CHS-like molecules were identified in the cryo-EM map (Extended data Fig. 11). These CHS molecules, along with other lipid-like densities, form a tightly packed “lipid belt” surrounding the TMD (Extended data Fig. 11a). We hypothesize that PregS may bind to one or more of these sites to activate TRPM3. Supporting this hypothesis, the gain-of-function mutation P1069Q, located adjacent to a CHS density, increases TRPM3 sensitivity and is associated with developmental and epileptic encephalopathies (Extended data Fig. 11b) ^4–6^. This suggests that CHS binding sites may overlap with or interact with PregS binding regions, influencing channel gating. Of note, a recent study proposed a putative PregS binding site that corresponds to one of the CHS binding sites identified in our study^50^ (Extended data Fig. 11, CHS site 3). However, given the moderate resolution and quality of the cryo-EM data in this study^50^, further research will be required to disentangle the effects of CHS and PregS and definitively identify PregS-mediated activation sites.

By elucidating the mechanisms of polymodal gating, this study establishes TRPM3 as a paradigm for understanding ion channel activation by diverse stimuli. The structural plasticity of TRPM3 underscores its ability to accommodate diverse modes of regulation. The identified ligand-binding sites, including the S1–S4 domain cavity and CHS binding sites, represent promising druggable targets. Furthermore, modulating the electrostatic interactions between MHR1/2 and the rib helix with small molecules could provide a novel approach to regulating TRPM3 activation. Together, these structural insights pave the way for the development of selective TRPM3 modulators to address critical unmet needs in pain management and neurological disorders.

---

**Extended Data Figure 1:**
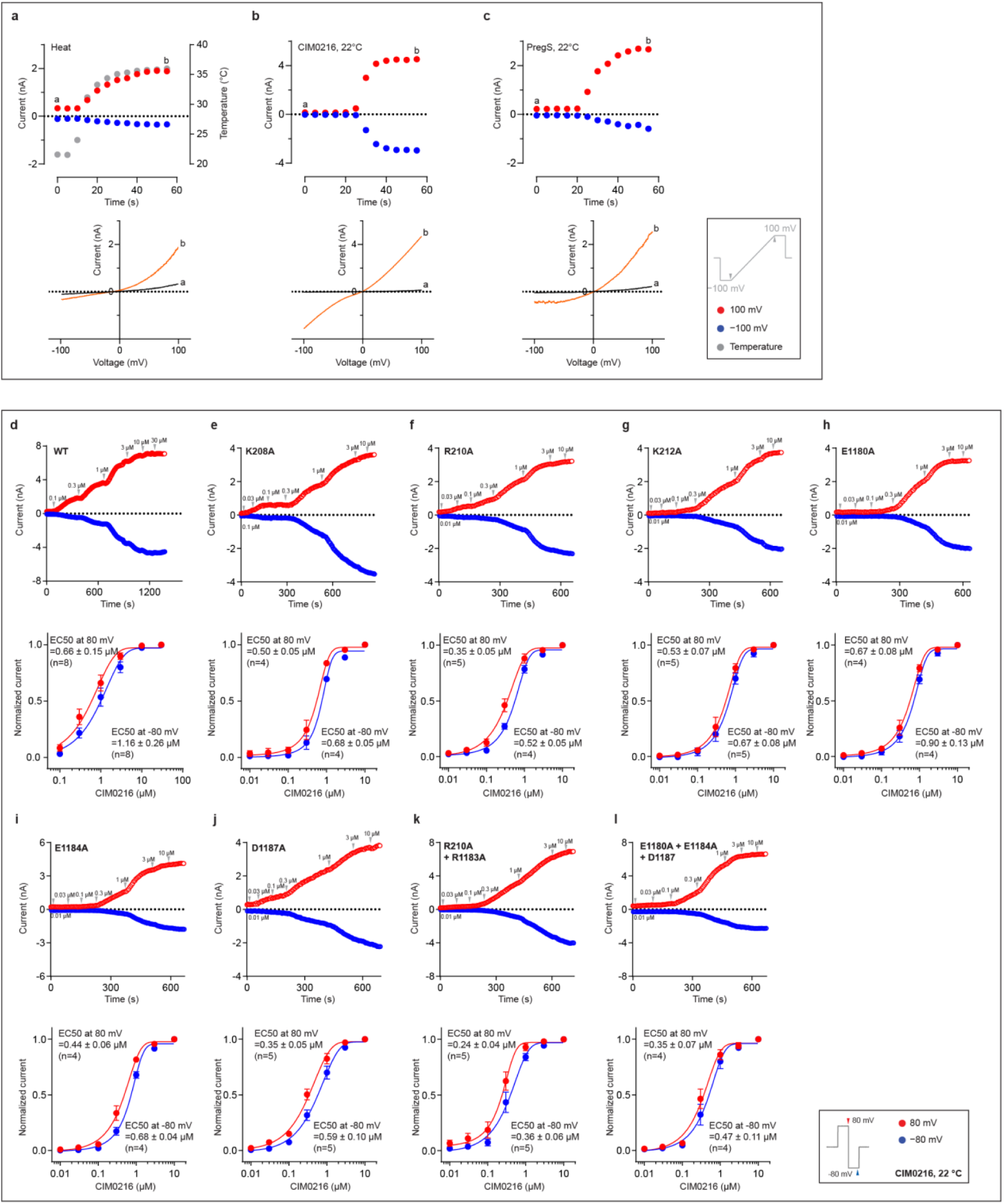
Electrophysiological characterization of TRPM3 in response to chemical and thermal stimulation. **a**, Representative whole-cell heat-activated currents measured in tsA cells overexpressing wild-type TRPM3. A voltage protocol was applied every 5 s to monitor current changes: −100 mV for 50 ms, ramped to +100 mV over 200 ms, and held at +100 mV for 50 ms. Currents were measured as the temperature increased from 22 °C to 37 °C (n = 5). Current amplitudes at +100 mV were plotted. The lower panel shows I/V curves corresponding to the time points indicated in the upper panel. **b**,**c**, Representative whole-cell currents measured in tsA cells overexpressing wild-type TRPM3 in response to CIM0216 (**b**, n = 5) and PregS (**c**, n = 5) using the same protocol as in panel (a). **d**–**l**, CIM0216 EC50 measurements for wild-type TRPM3. Upper panels show representative raw whole-cell currents, while lower panels present the fitted data. A protocol was applied every 5 s to monitor current changes: 0 mV for 50 ms, switch to +80 mV for 50 ms, followed −80 mV for 50 ms, and a return to 0 mV for 50 ms. Current amplitudes at +80 mV and −80 mV were plotted. The number of independent replicates (n) is indicated in each panel.

**Extended Data Figure 2:**
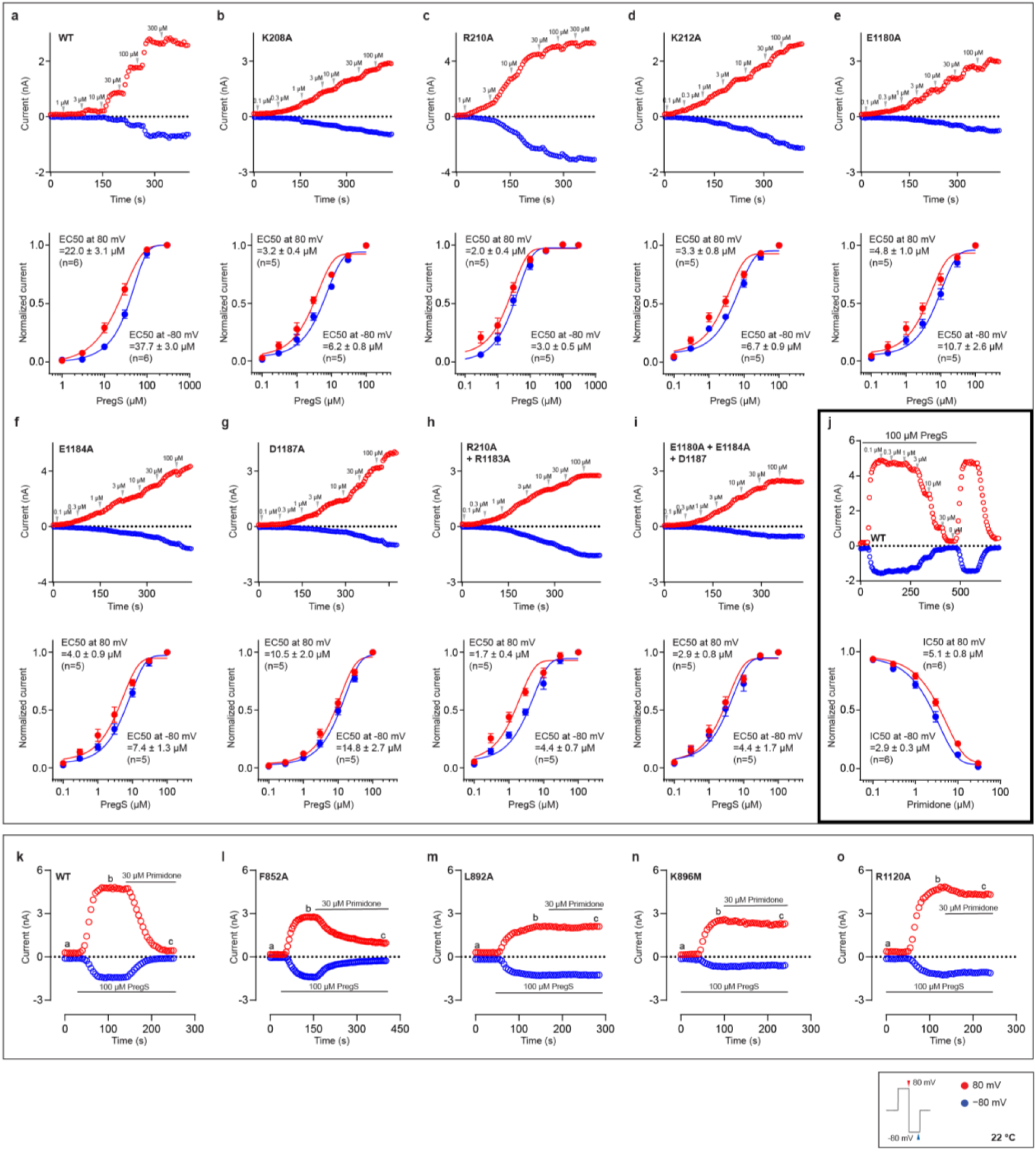
Electrophysiological characterization of TRPM3 in response to PregS and primidone. **a**–**i**, PregS EC50 measurements for wild-type TRPM3. Upper panels show representative raw whole-cell currents, while lower panels present the fitted data. A protocol was applied every 5 s to monitor current changes: 0 mV for 50 ms, switch to +80 mV for 50 ms, followed by −80 mV for 50 ms, and a return to 0 mV for 50 ms. Current amplitudes at +80 mV and −80 mV were plotted. The number of independent replicates (n) is indicated in each panel. **j**, Primidone IC50 measurements for wild-type TRPM3 induced by PregS using the same protocol as in panels (a–j). Upper panels show representative raw whole-cell currents, while lower panels present the fitted data. Current amplitudes at +80 mV and −80 mV were plotted. The number of independent replicates (n) is indicated in each panel. **k**–**o**, Primidone inhibition on PregS induced currents in wild-type TRPM3 and primidone binding site mutants, using the same protocol as in panels (a–j). Current amplitudes at +80 mV and −80 mV were plotted. The number of independent replicates (n) is indicated in each panel.

**Extended Data Figure 3:**
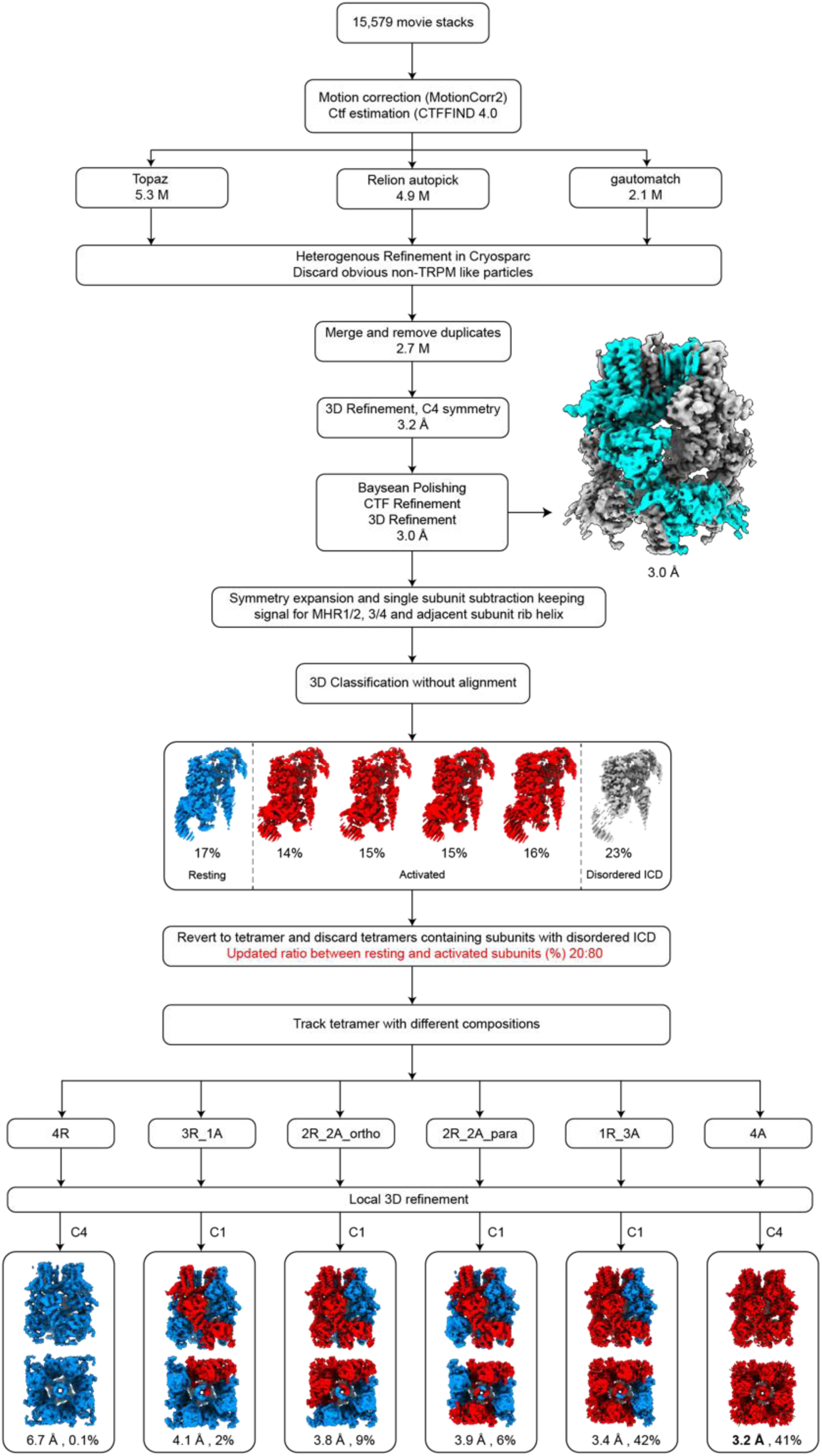
Cryo-EM data processing workflow of TRPM3 with CIM0216 at 18 °C. Key maps are shown along with the percentages of different subunit and tetramer conformations.

**Extended Data Figure 4:**
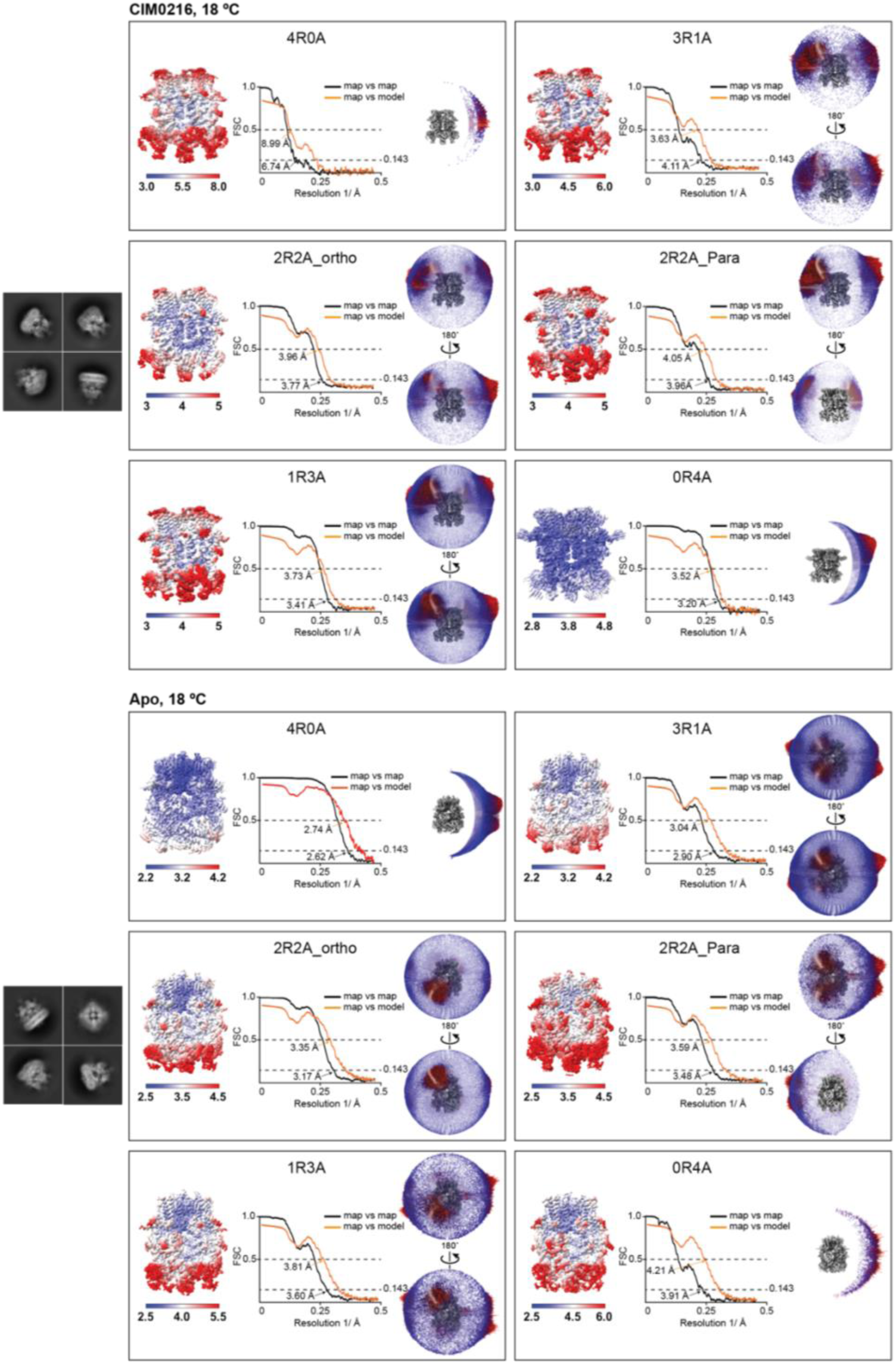
Cryo-EM analysis of TRPM3 bound to CIM0216 at 18 °C and TRPM3 in the apo state at 18 °C. Representative 2D class averages for each dataset are shown on the left. The labels 4R0A, 3R1A, 2R2A_ortho (orthogonal), 2R2A_para (parallel), 1R3A, and 0R4A denote tetrameric TRPM3 configurations composed of varying ratios of resting (R) and activated (A) subunits. For each tetrameric configuration, the local resolution estimation, FSC curves (map vs. map and map vs. model), and angular distribution of particles contributing to the final cryo-EM map reconstruction are displayed from left to right.

**Extended Data Figure 5:**
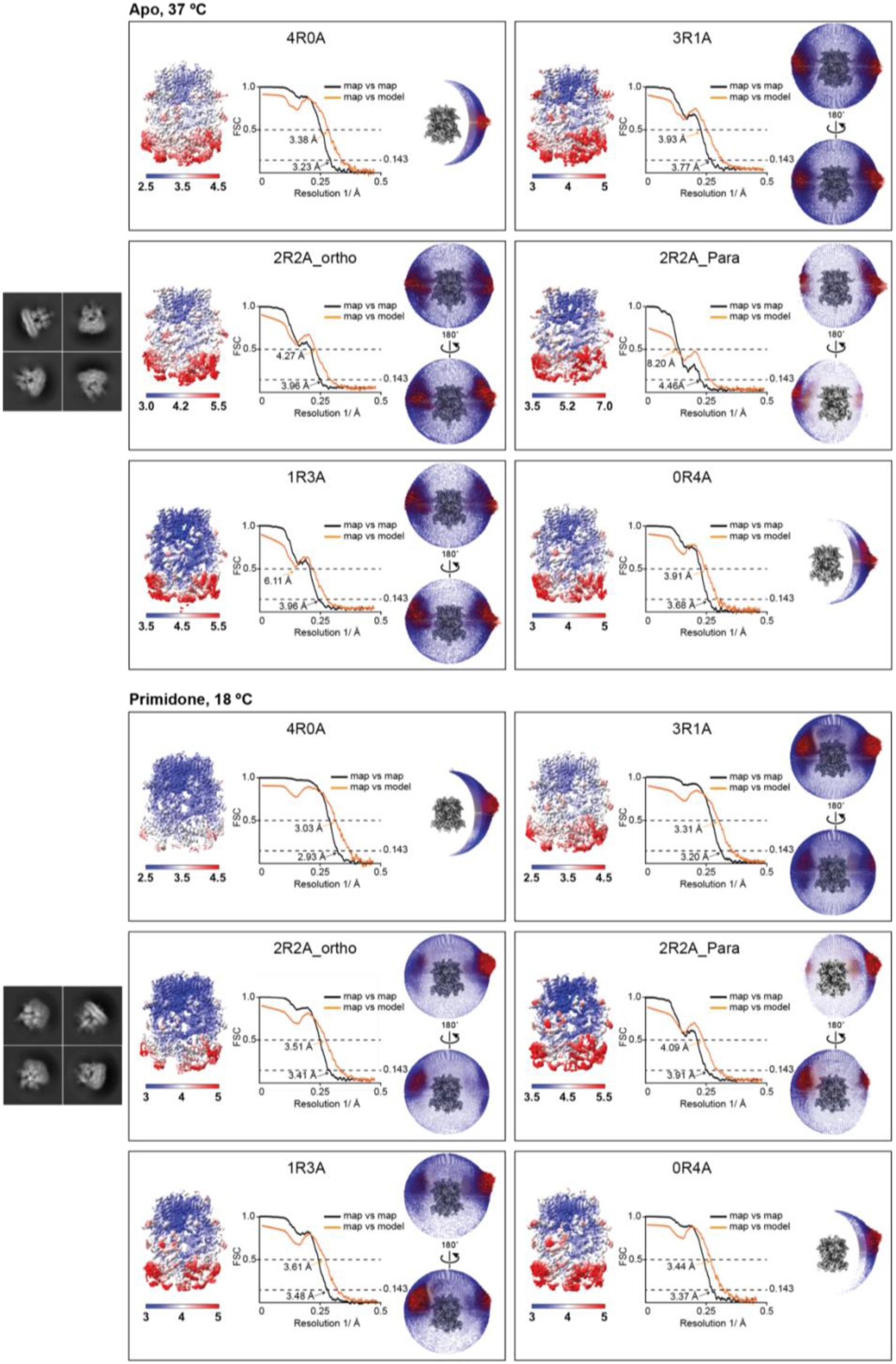
Cryo-EM analysis of TRPM3 in the apo state at 37 °C and TRPM3 bound to primidone at 18 °C. Representative 2D class averages for each dataset are shown on the left. The labels 4R0A, 3R1A, 2R2A_ortho (orthogonal), 2R2A_para (parallel), 1R3A, and 0R4A denote tetrameric TRPM3 configurations composed of varying ratios of resting (R) and activated (A) subunits. For each tetrameric configuration, the local resolution estimation, FSC curves (map vs. map and map vs. model), and angular distribution of particles contributing to the final cryo-EM map reconstruction are displayed from left to right.

**Extended Data Figure 6:**
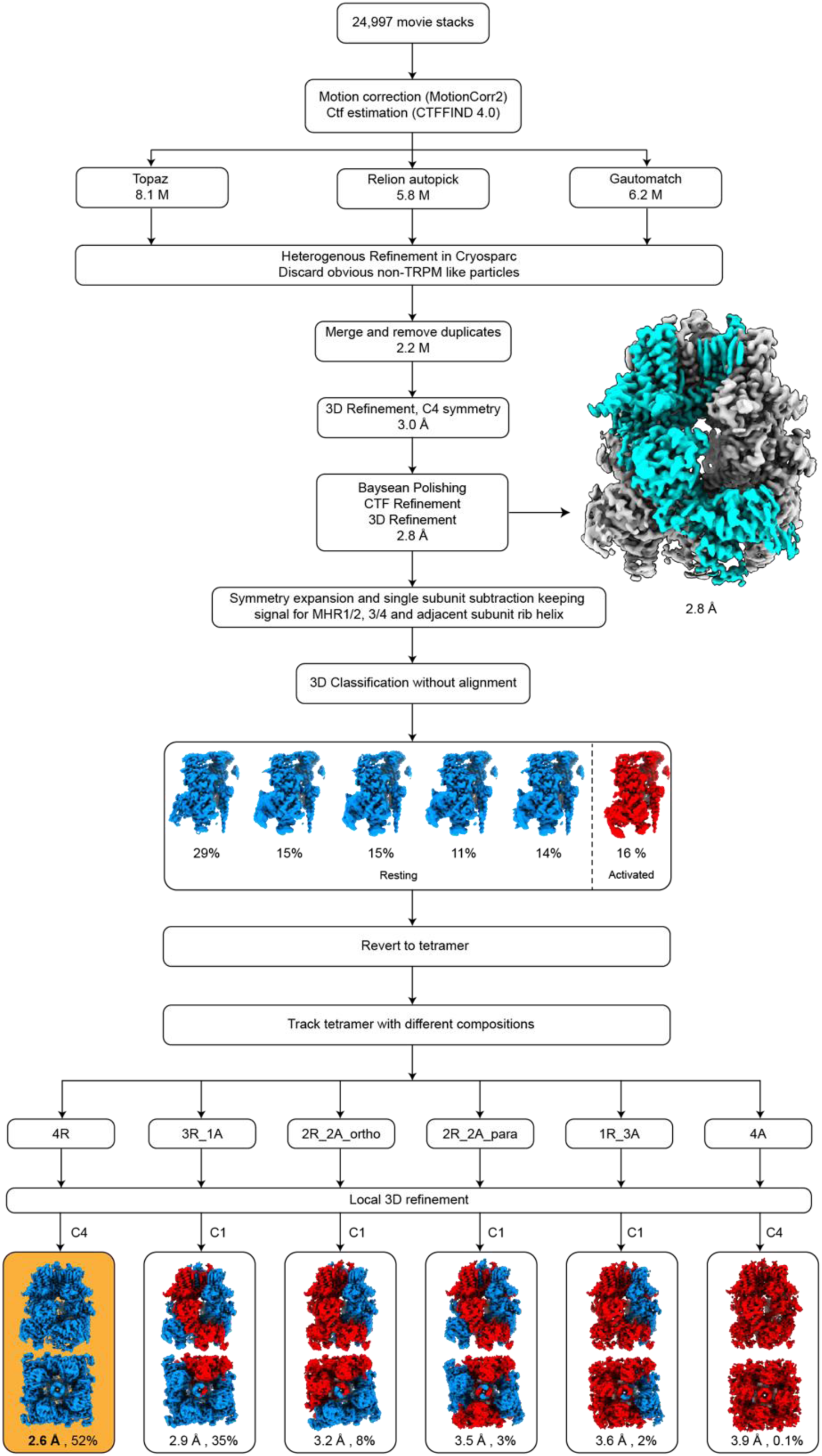
Cryo-EM data processing workflow of TRPM3 under apo condition at 18 °C. Key maps are shown along with the percentages of different subunit and tetramer conformations. The best tetramer map is highlighted with an orange background.

**Extended Data Figure 7:**
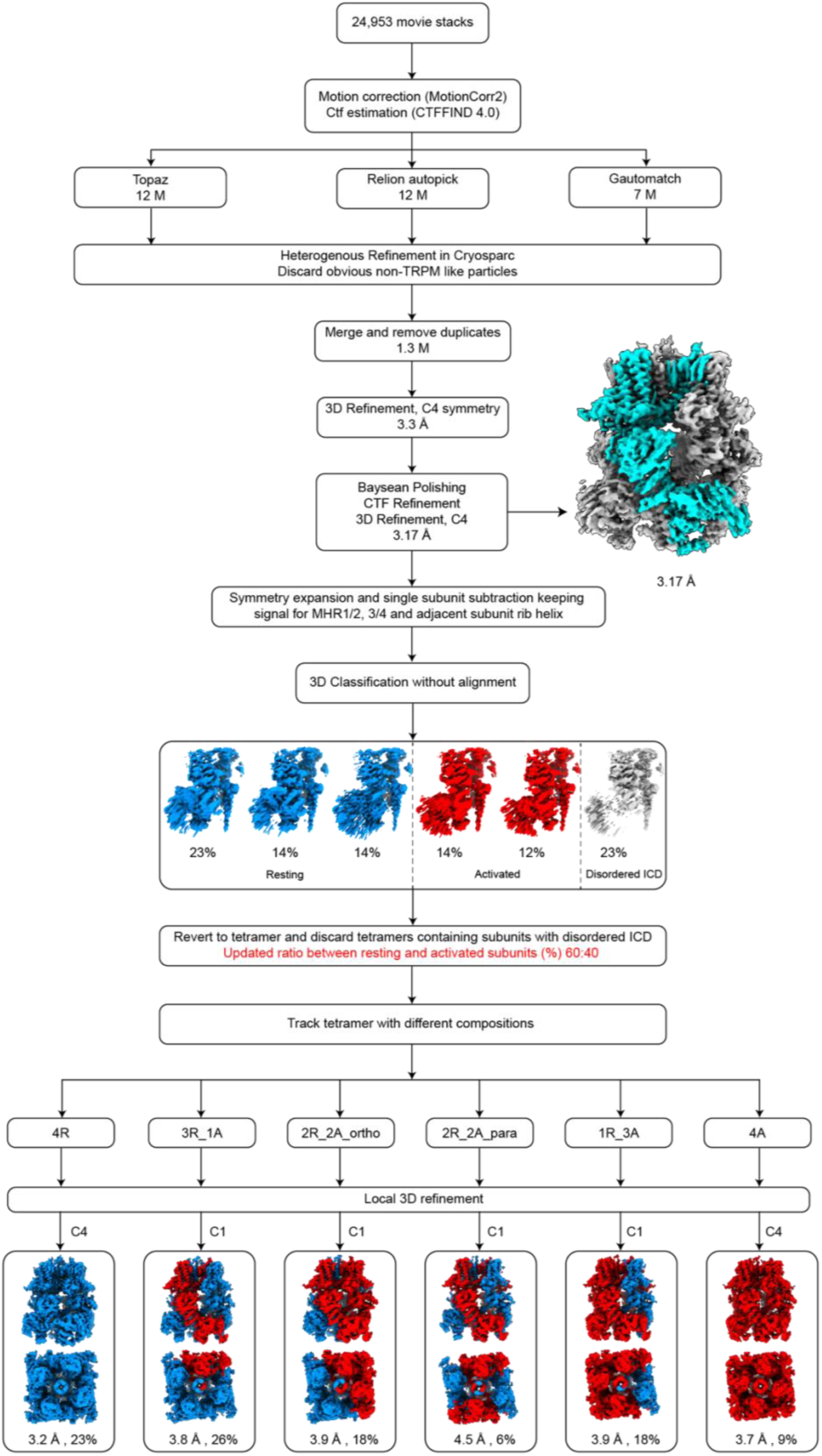
Cryo-EM data processing workflow of TRPM3 under apo condition at 37 °C. Key maps are shown along with the percentages of different subunit and tetramer conformations.

**Extended Data Figure 8:**
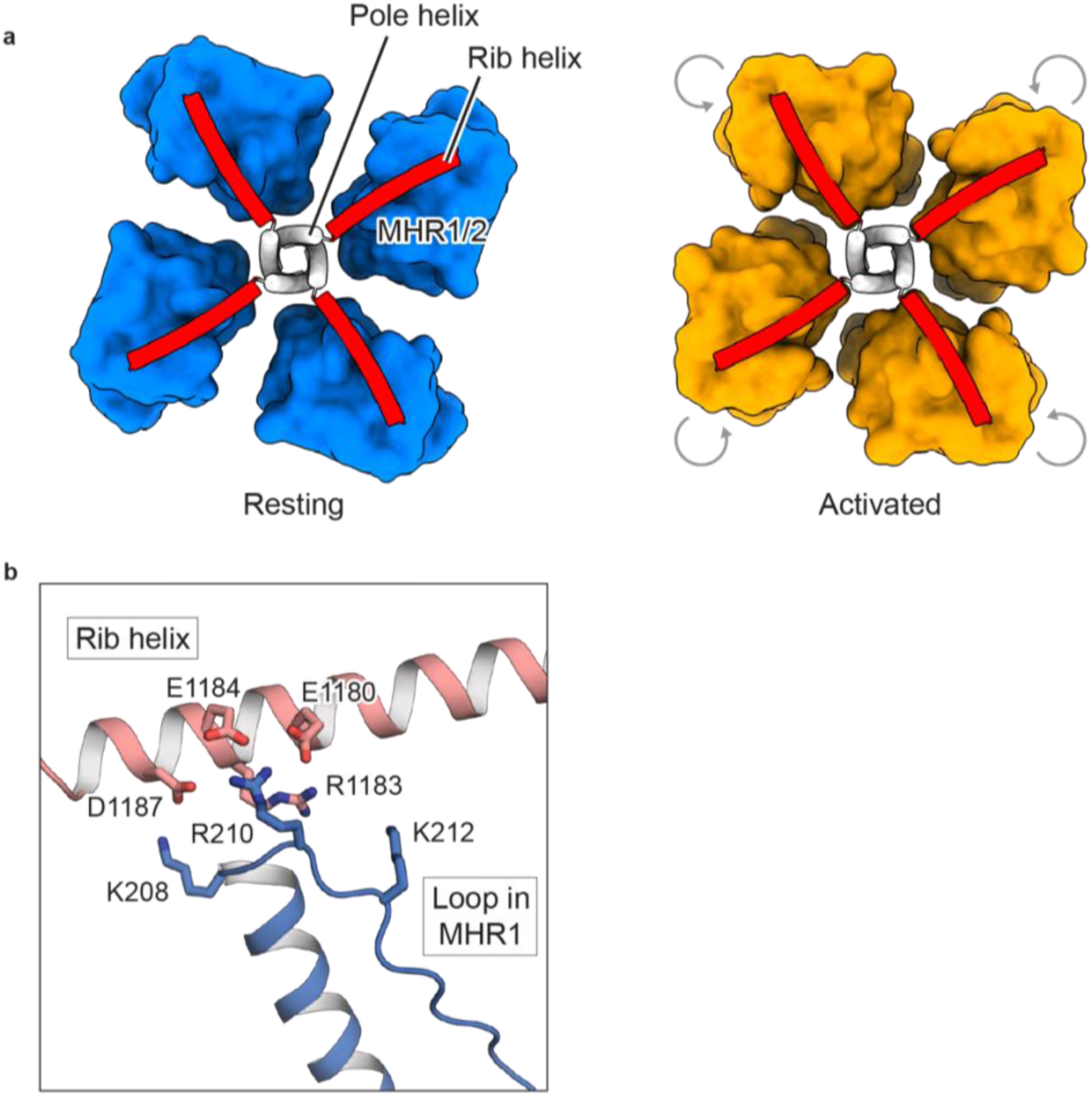
Conserved conformational changes in the ICD induced by heat or agonist binding. **a**, Both heat and agonist binding disrupt the interface between the MHR1/2 domain (depicted as blue and orange surfaces) and the adjacent rib helix (shown as a red cartoon). This disruption leads to a similar counterclockwise inward rotation of the MHR1/2 domain when viewed from the intracellular side. **b**, Close-up view of the interface in the resting state, highlighting the interaction between the positively charged loop in the MHR1/2 domain (blue) and the rib helix (red). Charged residues involved in these interactions are displayed in stick representation.

**Extended Data Figure 9:**
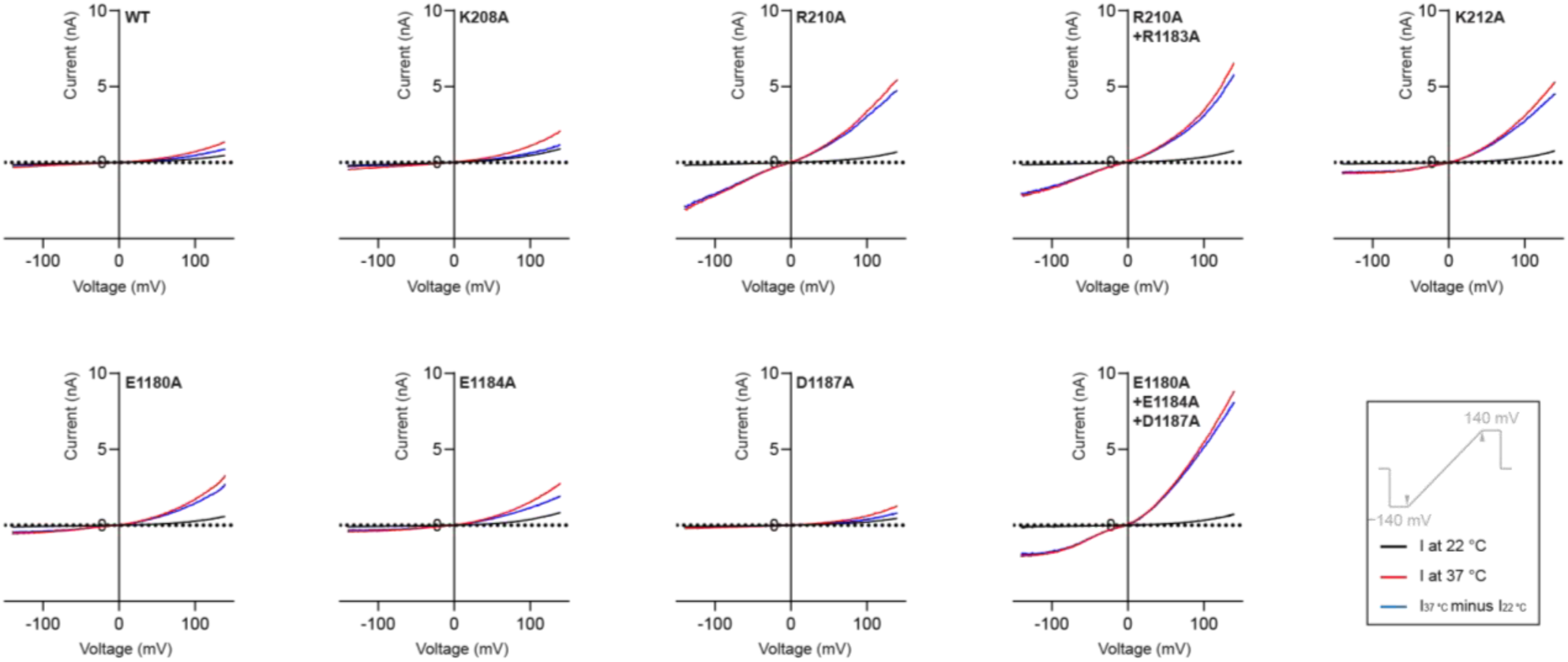
Mean whole-cell currents measured in tsA cells overexpressing wild-type TRPM3 or mutants at the interface between the MHR1/2 domain and the rib helix. A voltage protocol was applied every 5 s to monitor current changes: −140 mV for 50 ms, ramped to +140 mV over 200 ms, and held at +100 mV for 50 ms. Currents were measured as the temperature increased from 22 °C to 37 °C. Thermo currents at 140 mV were calculated by subtracting the steady-state current at 22 °C from the steady-state current at 37 °C. WT: n=9, K208A: n=5, R210A: n=7, R210A+R1183A: n=7, K212A: n=8, E1180A: n=8, E1184A: n=6, D1187A: n=5, E1180A+E1184A+D1187A: n=6.

**Extended Data Figure 10:**
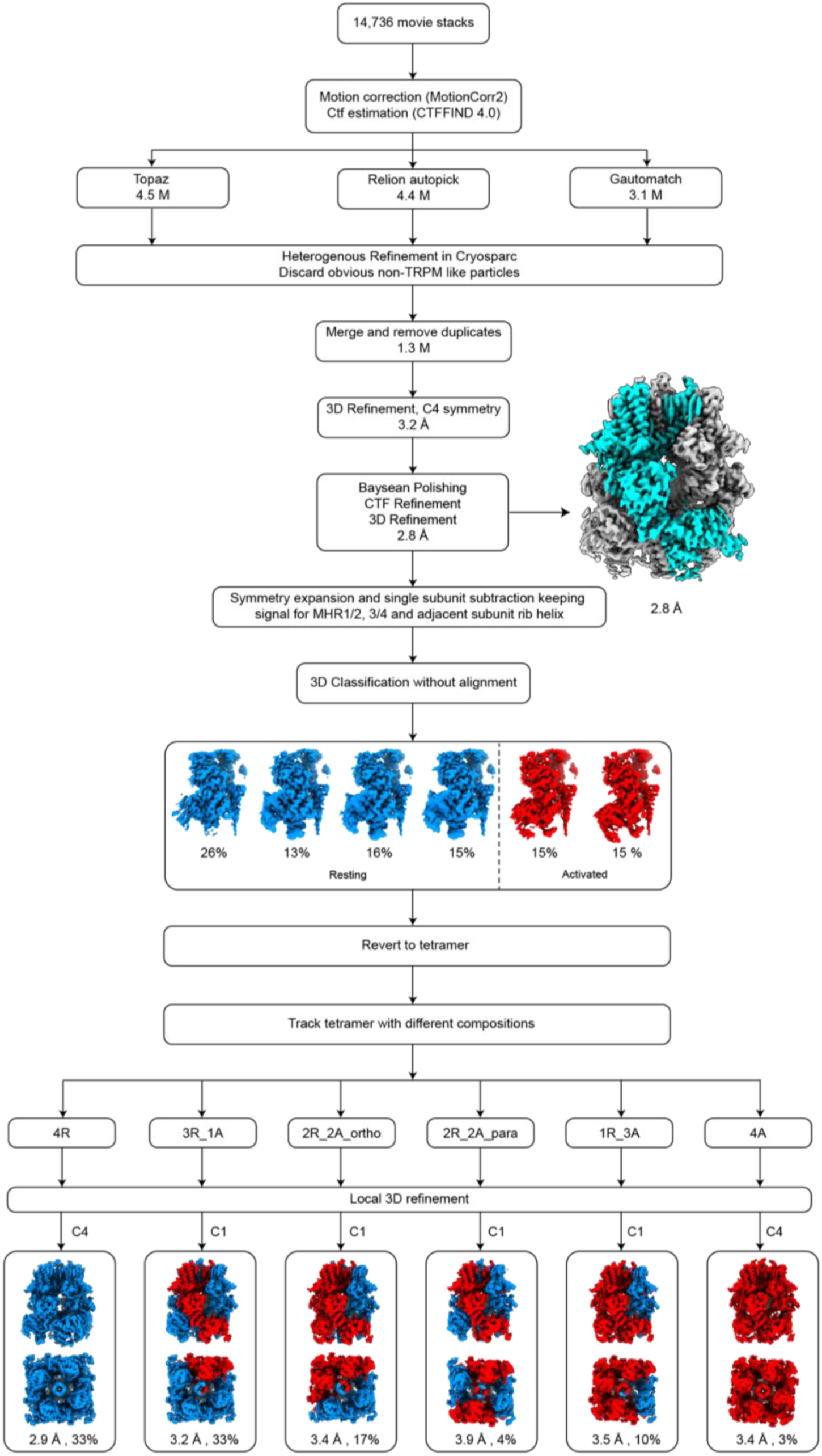
Cryo-EM data processing workflow of TRPM3 with primidone at 18 °C. Key maps are shown along with the percentages of different subunit and tetramer conformations.

**Extended Data Figure 11:**
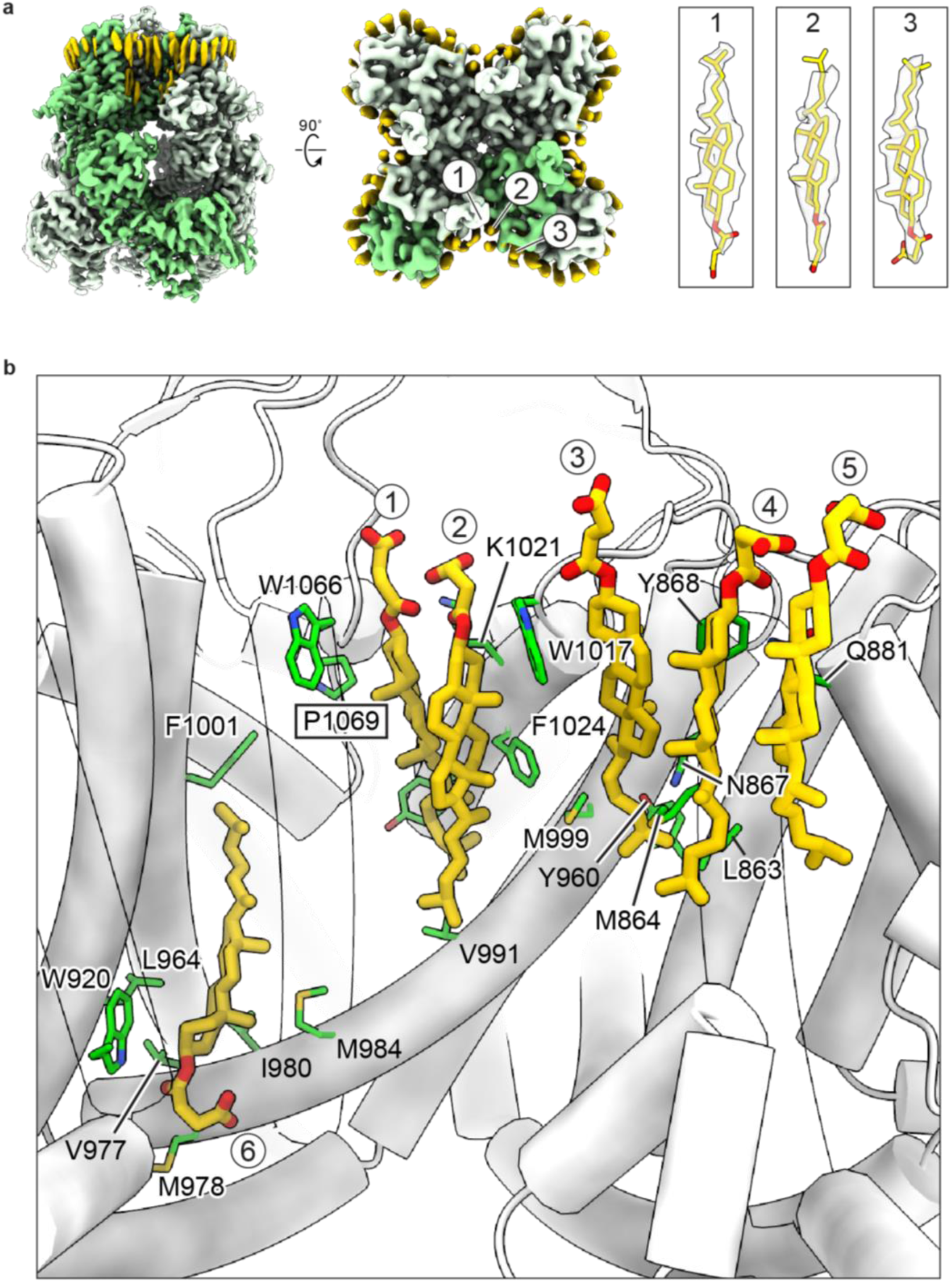
Bindings sites of cholesteryl hemisuccinate (CHS) **a**, Cryo-EM map of the apo resting state, highlighting one subunit in green and putative lipid densities in yellow, viewed parallel to the membrane (left) and from the intracellular side (middle). The three most well-defined CHS densities are labeled in the middle panel, with their corresponding densities displayed on the right panel. **b**, Close-up views of the CHS binding sites. The six CHS molecules are shown in yellow, with key interacting residues from the protein depicted in green. The position of the disease-associated mutation P1069Q is highlighted with a rectangle.

**Extended Data Table 1:** Cryo-EM data collection, refinement and validation statistics.

| CIM201, 18 °C |  |  |  |  |  |  |
| --- | --- | --- | --- | --- | --- | --- |
| Data Collection and Processing |  |  |  |  |  |  |
| Magnification | 105,000 |  |  |  |  |  |
| Voltage (kV) | 300 |  |  |  |  |  |
| Electron exposure (e-/Å<sup>2</sup>) | 58 |  |  |  |  |  |
| Defocus range (µm) | -1.1 to -1.4 |  |  |  |  |  |
| Pixel size (Å) | 0.826 |  |  |  |  |  |
| Initial particle images (no.) | 2,662,548 |  |  |  |  |  |
| Maps | 4R | 3R1A | 2R2A_ortho | 2R2A_para | 1R3A | 0R4A |
| Symmetry imposed | C4 | C1 | C1 | C1 | C1 | C4 |
| Final particle images (no.) | 1642 | 29177 | 127753 | 81343 | 577875 | 563237 |
| Map resolution (Å) | 6.74 | 4.11 | 3.77 | 3.96 | 3.41 | 3.20 |
| FSC threshold | 0.143 | 0.143 | 0.143 | 0.143 | 0.143 | 0.143 |
| Map resolution range (Å) | 4.47-19.21 | 3.84-14.48 | 3.43-10.75 | 3.48-12.19 | 3.20-8.01 | 3.03-6.28 |
| Refinement |  |  |  |  |  |  |
| Initial model used (PDB code) | 8DDQ |  |  |  |  | 8DDQ |
| Model resolution (Å) | 8.99 |  |  |  |  | 3.52 |
| FSC threshold | 0.5 |  |  |  |  | 0.5 |
| Model resolution range (Å) | 8.99-91.74 |  |  |  |  | 3.52-91.74 |
| Map sharpening *B* factor (Å<sup>2</sup>) | -93.94 |  |  |  |  | -114.18 |
| Model composition |  |  |  |  |  |  |
| Non-hydrogen atoms | 30976 |  |  |  |  | 30808 |
| Protein residues | 4084 |  |  |  |  | 4068 |
| Ligands | 4 |  |  |  |  | 4 |
| *B* factors (Å<sup>2</sup>) |  |  |  |  |  |  |
| Protein | 273.52 |  |  |  |  | 139.82 |
| Ligand | 225.46 |  |  |  |  | 119.37 |
| R.m.s. deviations |  |  |  |  |  |  |
| Bond lengths (Å) | 0.004 |  |  |  |  | 0.004 |
| Bond angles (°) | 0.769 |  |  |  |  | 0.793 |
| Validation |  |  |  |  |  |  |
| MolProbity score | 2.24 |  |  |  |  | 2.33 |
| Clashscore | 27.30 |  |  |  |  | 18.44 |
| Poor rotamers (%) | 0 |  |  |  |  | 1.23 |
| Ramachandran plot |  |  |  |  |  |  |
| Favored (%) | 95.41 |  |  |  |  | 93.82 |
| Allowed (%) | 4.49 |  |  |  |  | 6.18 |
| Disallowed (%) | 0.1 |  |  |  |  | 0 |

**Extended Data Table 2:** Cryo-EM data collection, refinement and validation statistics.

| Apo, 18 °C |  |  |  |  |  |  |
| --- | --- | --- | --- | --- | --- | --- |
| Data Collection and Processing |  |  |  |  |  |  |
| Magnification | 105,000 |  |  |  |  |  |
| Voltage (kV) | 300 |  |  |  |  |  |
| Electron exposure (e-/Å<sup>2</sup>) | 58 |  |  |  |  |  |
| Defocus range (µm) | -1.1 to -1.4 |  |  |  |  |  |
| Pixel size (Å) | 0.826 |  |  |  |  |  |
| Initial particle images (no.) | 2,191,867 |  |  |  |  |  |
| Maps | 4R | 3R1A | 2R2A_ortho | 2R2A_para | 1R3A | 0R4A |
| Symmetry imposed | C4 | C1 | C1 | C1 | C1 | C4 |
| Final particle images (no.) | 1138501 | 762853 | 187643 | 62369 | 37679 | 2822 |
| Map resolution (Å) | 2.62 | 2.90 | 3.17 | 3.48 | 3.60 | 3.91 |
| FSC threshold | 0.143 | 0.143 | 0.143 | 0.143 | 0.143 | 0.143 |
| Map resolution range (Å) | 2.35-5.30 | 2.60-7.08 | 2.88-9.0 | 3.12-11.88 | 4.77-12.21 | 3.40-13.45 |
| Refinement |  |  |  |  |  |  |
| Initial model used (PDB code) | 8DDQ |  |  |  |  | 8DDQ |
| Model resolution (Å) | 2.74 |  |  |  |  | 4.21 |
| FSC threshold | 0.5 |  |  |  |  | 0.5 |
| Model resolution range (Å) | 2.74-91.74 |  |  |  |  | 4.21-91.74 |
| Map sharpening *B* factor (Å<sup>2</sup>) | -88.05 |  |  |  |  | -43.32 |
| Model composition |  |  |  |  |  |  |
| Non-hydrogen atoms | 33136 |  |  |  |  | 32704 |
| Protein residues | 4272 |  |  |  |  | 4216 |
| Ligands | 0 |  |  |  |  | 0 |
| *B* factors (Å<sup>2</sup>) |  |  |  |  |  |  |
| Protein | 126.91 |  |  |  |  | 160.34 |
| Ligand | 0 |  |  |  |  | 0 |
| R.m.s. deviations |  |  |  |  |  |  |
| Bond lengths (Å) | 0.003 |  |  |  |  | 0.004 |
| Bond angles (°) | 0.690 |  |  |  |  | 0.913 |
| Validation |  |  |  |  |  |  |
| MolProbity score | 1.30 |  |  |  |  | 1.63 |
| Clashscore | 3.22 |  |  |  |  | 5.99 |
| Poor rotamers (%) | 0.79 |  |  |  |  | 0.80 |
| Ramachandran plot |  |  |  |  |  |  |
| Favored (%) | 96.88 |  |  |  |  | 95.63 |
| Allowed (%) | 3.03 |  |  |  |  | 4.17 |
| Disallowed (%) | 0.09 |  |  |  |  | 0.19 |

**Extended Data Table 3:** Cryo-EM data collection, refinement and validation statistics.

| Apo, 37 °C |  |  |  |  |  |  |
| --- | --- | --- | --- | --- | --- | --- |
| Data Collection and Processing |  |  |  |  |  |  |
| Magnification | 105,000 |  |  |  |  |  |
| Voltage (kV) | 300 |  |  |  |  |  |
| Electron exposure (e-/Å²) | 58 |  |  |  |  |  |
| Defocus range (µm) | -1.1 to -1.4 |  |  |  |  |  |
| Pixel size (Å) | 0.826 |  |  |  |  |  |
| Initial particle images (no.) | 1,252,197 |  |  |  |  |  |
| Maps | 4R | 3R1A | 2R2A_ortho | 2R2A_para | 1R3A | 0R4A |
| Symmetry imposed | C4 | C1 | C1 | C1 | C1 | C4 |
| Final particle images (no.) | 111874 | 125354 | 84256 | 30726 | 84811 | 40005 |
| Map resolution (Å) | 3.23 | 3.77 | 3.96 | 4.46 | 3.96 | 3.68 |
| FSC threshold | 0.143 | 0.143 | 0.143 | 0.143 | 0.143 | 0.143 |
| Map resolution range (Å) | 2.96-7.73 | 3.22-9.92 | 3.43-12.51 | 3.87-16.89 | 3.54-11.77 | 3.25-8.76 |
| Refinement |  |  |  |  |  |  |
| Initial model used<br>(PDB code) | 8DDQ |  |  |  |  | 8DDQ |
| Model resolution (Å) | 3.38 |  |  |  |  | 3.91 |
| FSC threshold | 0.5 |  |  |  |  | 0.5 |
| Model resolution range (Å) | 3.38-91.74 |  |  |  |  | 3.91-91.74 |
| Map sharpening *B* factor (Å²) | -95.17 |  |  |  |  | -95.84 |
| Model composition |  |  |  |  |  |  |
| Non-hydrogen atoms | 34576 |  |  |  |  | 33024 |
| Protein residues | 4220 |  |  |  |  | 4146 |
| Ligands | 0 |  |  |  |  | 0 |
| *B* factors (Å²) |  |  |  |  |  |  |
| Protein | 135.73 |  |  |  |  | 143.77 |
| Ligand | 0 |  |  |  |  | 0 |
| R.m.s. deviations |  |  |  |  |  |  |
| Bond lengths (Å) | 0.003 |  |  |  |  | 0.005 |
| Bond angles (°) | 0.597 |  |  |  |  | 0.919 |
| Validation |  |  |  |  |  |  |
| MolProbity score | 2.03 |  |  |  |  | 2.94 |
| Clashscore | 19.48 |  |  |  |  | 26.79 |
| Poor rotamers (%) | 0.85 |  |  |  |  | 1.84 |
| Ramachandran plot |  |  |  |  |  |  |
| Favored (%) | 96.37 |  |  |  |  | 93.05 |
| Allowed (%) | 3.63 |  |  |  |  | 6.76 |
| Disallowed (%) | 0 |  |  |  |  | 0.2 |

**Extended Data Table 4:** Cryo-EM data collection, refinement and validation statistics.

| Primidone, 18 °C |  |  |  |  |  |  |
| --- | --- | --- | --- | --- | --- | --- |
| Data Collection and Processing |  |  |  |  |  |  |
| Magnification | 105,000 |  |  |  |  |  |
| Voltage (kV) | 300 |  |  |  |  |  |
| Electron exposure (e-/Å²) | 58 |  |  |  |  |  |
| Defocus range (µm) | -1.1 to -1.4 |  |  |  |  |  |
| Pixel size (Å) | 0.826 |  |  |  |  |  |
| Initial particle images (no.) | 1,271,284 |  |  |  |  |  |
| Maps | 4R | 3R1A | 2R2A_ortho | 2R2A_para | 1R3A | 0R4A |
| Symmetry imposed | C4 | C1 | C1 | C1 | C1 | C4 |
| Final particle images (no.) | 419231 | 415433 | 213123 | 56912 | 122730 | 43855 |
| Map resolution (Å) | 2.93 | 3.20 | 3.41 | 3.91 | 3.48 | 3.37 |
| FSC threshold | 0.143 | 0.143 | 0.143 | 0.143 | 0.143 | 0.143 |
| Map resolution range (Å) | 2.68-6.43 | 2.90-9.10 | 3.02-11.08 | 3.38-13.23 | 3.12-12.13 | 3.01-12.01 |
| Refinement |  |  |  |  |  |  |
| Initial model used (PDB code) | 8DDQ |  |  |  |  | 8DDQ |
| Model resolution (Å) | 3.03 |  |  |  |  | 3.44 |
| FSC threshold | 0.5 |  |  |  |  | 0.5 |
| Model resolution range (Å) | 3.03-91.74 |  |  |  |  | 3.44-91.74 |
| Map sharpening *B* factor (Å²) | -96.03 |  |  |  |  | -92.95 |
| Model composition |  |  |  |  |  |  |
| Non-hydrogen atoms | 33046 |  |  |  |  | 32648 |
| Protein residues | 4240 |  |  |  |  | 4212 |
| Ligands | 4 |  |  |  |  | 4 |
| *B* factors (Å²) |  |  |  |  |  |  |
| Protein | 147.88 |  |  |  |  | 202.79 |
| Ligand | 96.19 |  |  |  |  | 104.71 |
| R.m.s. deviations |  |  |  |  |  |  |
| Bond lengths (Å) | 0.004 |  |  |  |  | 0.004 |
| Bond angles (°) | 0.848 |  |  |  |  | 0.874 |
| Validation |  |  |  |  |  |  |
| MolProbity score | 2.18 |  |  |  |  | 2.22 |
| Clashscore | 16.60 |  |  |  |  | 20.98 |
| Poor rotamers (%) | 1.15 |  |  |  |  | 0.80 |
| Ramachandran plot |  |  |  |  |  |  |
| Favored (%) | 95.51 |  |  |  |  | 94.71 |
| Allowed (%) | 4.30 |  |  |  |  | 5.10 |
| Disallowed (%) | 0.19 |  |  |  |  | 0.19 |

## Methods

### Constructs and mutagenesis

The full-length rabbit TRPM3 (rTRPM3) gene (Uniprot G1T6W8_RABIT) was subcloned into pEG BacMam vector with an N-terminal 8XHis-Twin-Strep-Alfa followed by a thrombin cleavage site^51^. Site-directed mutagenesis was performed by QuickChange site-directed mutagenesis protocol (Agilent) and confirmed by sequencing (Eurofins or Plasmidsaurus).

### TRPM3 protein expression and purification

For bacmid generation, pEG BacMam vector was transformed into DH10Bac cells. SF9 cells were transfected with freshly prepared bacmids for generation of P1 and subsequently P2 baculovirus. P2 viruses (8%) were used to infect tSA201 cells grown in Freestyle 293 expression medium (Gibco) in suspension culture. The cells were initially incubated at 37 °C for about 10–12 h. Thereafter, sodium butyrate was added to a final concentration of 10 mM and temperature was reduced to 30 °C to boost expression. The cells were harvested after a total growth of about 65 h, washed with buffer containing 20 mM Hepes pH 7.5 and 150 mM NaCl (HBS) and pelleted by centrifugation. The cell pellets were either processed immediately or flash frozen in Liq. N2 and stored at –80 °C.

The entire protein purification was carried out at 4 °C. The cells were suspended in HBS buffer containing 10 mM β-ME, 1 mM phenylmethylsulphonyl fluoride, 0.8 μM aprotinin, 2 μg/ml leupeptin and 2 mM pepstatinA and lysed by sonication. The cell lysate was centrifuged at 8000g for 20 mins to remove cell debris. The membrane fractions were collected by centrifugation at 186,000g for 1 h in an ultracentrifuge (Beckman Coulter). The membranes were resuspended in HBS containing 10mM β-ME, 2mM Mg^2+^/ATP, and protease inhibitors and homogenized using a Dounce homogenizer. After resuspension, membranes were solubilized in 1% LMNG/CHS (5:1 wt/wt) for about 1.5 hours at 4 °C followed by ultracentrifugation at 186,000g for 1 hour. The supernatant was loaded on to Strep-Tactin Superflow resin (IBA) at a flow rate of 0.1 ml/min. Thereafter, the resin was washed twice. The first wash was given with HBS buffer containing 0.01 % LMNG/CHS (5:1 wt/wt), 10 mM β-ME and 1mM Mg^2+^/ATP followed by a second wash with HBS buffer containing 0.003 % LMNG/CHS (5:1 wt/wt), 10mM β-ME and 5 mM EDTA. The protein was eluted with HBS buffer containing 0.003 % LMNG/CHS (5:1 wt/wt), 10 mM β-ME, 5 mM EDTA, and 10 mM Desthiobiotin. The elution was concentrated to 0.5 ml and further purified by size exclusion chromatography using Superose 6 increase 10/300 GL column (GE Healthcare) in a HBS buffer having 0.001 % LMNG/CHS (5:1 wt/wt), 10mM β-ME and 5 mM EDTA. The peak SEC fractions were pooled and concentrated to about 8 mg/mL for freezing cryo-EM grids.

### Cryo-EM sample preparation and data acquisition

Grids for the apo condition were prepared both at 18 and 37 °C. 2.5 ul of protein sample was applied to a freshly glow-discharged Quantifoil holey carbon grid (Au 1.2/1.3, 300 mesh) and after a wait time of 15 s at 37 °C, the grids were blotted for 1.5s in the Vitrobot Mark III set to 100 % humidity and a blot force of −15. The grids were plunged frozen into liquid ethane cooled by liquid nitrogen. For sample preparation at 18 °C, 2.5 ul of protein sample was applied to a freshly glow-discharged Quantifoil holey carbon grid (Au 1.2/1.3, 300 mesh). After waiting for 5 s, the grids were blotted for 1s in the Vitrobot Mark III set to 18 °C, 100 % humidity, and a blot force of 0. For CIM and Primidone containing conditions, the purified protein was mixed with 1mM CIM or Primidone and incubated for 1h. Thereafter, grids were frozen in the same manner as described for the apo condition at 18 °C. The cryoEM data were collected using FEI Titan Krios electron microscope operating at 300kV and a nominal magnification of 105,000X and an energy filter (20 eV slit width). Movies were recorded in super-resolution mode using a Gatan K3 Summit direct electron detector with a binned pixel size of 1.16 Angstrom. Nominal defocus ranged from –1.4 to –1.1 μm with a total dose of 58 e^-^.

### Cryo-EM data processing and analysis

Super-resolution movie stacks were motion corrected and 2X binned using MotionCorr2 (v.1.1.0)^52^. The contrast transfer function (CTF) parameters were estimated by ctffind (v.4.1.10)^53^. Particles were picked by Relion’s template-picking^54^, Gautomatch (v.0.56) (https://github.com/JackZhang-Lab/Gautmatch), and Topaz (v.0.2.4)^55^. Each particle set was independently cleaned by removing only obviously non-TRPM3-like junk particles by rounds of heterogeneous refinement in Cryosparc^56^. The remaining particles from the three cleanups were merged and duplicates removed. The deduplicated particles were used for homogenous refinement and non-uniform refinement with C4 symmetry in Cryosparc. The particles were further subjected to multiple rounds of CTF-refinement and Baysean polishing in Relion followed by 3D refinement with C4 symmetry to improve the map resolution.

The resulting particles were symmetry-expanded at the single subunit level with C4 symmetry. Using a mask encompassing single subunit ICD and rib-helix from the neighboring subunit, signal subtraction was carried out. The subtracted images were used for 3D classification without image alignment in Relion. The tetrameric particles containing broken/disordered ICD were excluded from further analysis, as they likely represent damaged particles during cryo-EM sample preparation. The remaining intact tetrameric particles were pooled for subsequent analysis. The ratio of subunits in the resting and activated states within these intact tetrameric particles was calculated. Subsequently, subunit particles in the resting and activated states were traced back to their respective tetrameric assemblies, resulting in six distinct compositions: 4 resting and 0 activated (4R0A), 3 resting and 1 activated (3R1A), 2 resting and 2 activated in orthogonal positions (2R2A_ortho), 2 resting and 2 activated in parallel positions (2R2A_para), 1 resting and 3 activated (1R3A), and 0 resting and 4 activated (0R4A) particles. The two homotetrameric compositions (4R0A and 0R4A) underwent further local refinement with C4 symmetry. For the heterotetrameric compositions, particles were rotated to align the positions of resting and activated subunits within the tetramer, ensuring proper orientation, and subsequently underwent local refinement with C1 symmetry in Relion.

The detailed image processing steps are summarized and illustrated in Extended Data Figs 3, 6, 7 and 10. Map resolution estimates were based on the gold standard Fourier shell correlation (FSC) 0.143 criteria for all the datasets.

### Model building

For building atomic models of rTRPM3, the published mouse TRPM3 structure (PDB-ID 8DDQ) was docked into the final cryoEM map by rigid-body fitting. Thereafter, the residues were mutated to rTRPM3 and manually fit into the map using Coot. For obtaining the flipped subunit conformation, the MHR domains, TMD, and C-terminal domains of the non-flipped conformations were fitted into the flipped-conformation map by rigid-body fitting and then manually adjusted in Coot^57^. Ligands were manually fitted into the density in Coot and refined in real space. All the models were subjected to real space refinement in Phenix^58^ to improve the model metrices. The final models were validated using MolProbity^59^. All the figures were prepared using UCSF ChimeraX^60^ and PyMOL (Schrodinger LLC).

### Electrophysiology

TsA201 cells expressing plasmids encoding N-terminal GFP tagged rabbit TRPM3 were used. After one day post-transfection with plasmid DNA (100 ng/mL) and Lipofectamine 2000 (Invitrogen, 11668019), the cells were trypsinized and replated onto poly-L-lysine-coated (Sigma P4707) glass coverslips After cell attachment, the coverslip was transferred to a recording chamber. Whole-cell patch-clamp recordings were performed at room temperature (21–23°C) or body temperature (36– 38°C). The temperature of perfusion solutions was controlled by thermal control devices (SC-20/CL-100, Warner Instruments, Holliston, MA).

Signals were amplified using a Multiclamp 700B amplifier and digitized using a Digidata 1550B A/D converter (Molecular Devices, Sunnyvale CA). The whole-cell current was measured on the cells with an access resistance of less than 10 MΩ after the whole-cell configuration was obtained. The whole-cell capacitance was compensated by the amplifier circuitry. The ramp pulse from −140 to 140 mV (or from –100 to 100 mV) for 200 ms or the two-step pulse from 80 to –80 mV for 50 ms was continuously applied to the cell membrane every 5 sec to monitor the TRPM3 current. Electrical signals were digitized at 10 kHz and filtered at 2 kHz. Recordings were analyzed using Clampfit 11.3 (Axon Instruments Inc), GraphPad Prism 10 (La Jolla, CA), and OriginPro 2024 (OriginLab, Northampton, MA). The standard bath solution contains (in mM): 150 NaCl, 5 KCl, 1 MgCl_2_, 2 CaCl_2_, 12 Mannitol, 10 HEPES, pH=7.4 with NaOH. For a whole cell recording, the extracellular and intracellular solutions contain (in mM): 150 NaCl, 10 HEPES, 1 MgCl_2_, 5 EGTA.

## Data availability

Cryo-EM density maps have been deposited at the EMDB (Electron Microscopy Data Bank) and the Research Collaboratory for Structural Bioinformatics Protein Data Bank (RCS-PDB), respectively.

## Acknowledgements

We thank G. Zhao and X. Meng for the support with data collection at the David Van Andel Advanced Cryo-Electron Microscopy Suite. We appreciate the high-performance computing team of VAI, structural biology facility and Quest High-Performance Computing Cluster at Northwestern University for computational support. W.L. is supported by National Institutes of Health (NIH) grants (R01HL153219 and R01NS112363). J.D. is supported by a McKnight Scholar Award, a Klingenstein-Simon Scholar Award, a Sloan Research Fellowship in neuroscience, a Pew Scholar in the Biomedical Sciences award, and NIH grants (R01NS111031 and R01NS129804).

## Author Contributions

J.D. and W.L. supervised the project. W.C., S.V. and S.K. initiated the project. S.K. and F.J. carried out protein purification, cryo-EM data collection, and processing. R.M. and S.K. generated the mutants and carried out construct screening. S.P. performed electrophysiological experiments. J.D., W.L., F.J., S.K., and S.P. contributed to the manuscript preparation. The authors declare no conflicts of interest.

## References

1 Vriens, J., Nilius, B. & Voets, T. Peripheral thermosensation in mammals. Nat Rev Neurosci 15, 573–589 (2014). 10.1038/nrn3784

2 Basbaum, A. I., Bautista, D. M., Scherrer, G. & Julius, D. Cellular and molecular mechanisms of pain. Cell 139, 267–284 (2009). 10.1016/j.cell.2009.09.028

3 Lee, N. et al. Expression and characterization of human transient receptor potential melastatin 3 (hTRPM3). J Biol Chem 278, 20890–20897 (2003). 10.1074/jbc.M211232200

4 Van Hoeymissen, E. et al. Gain of channel function and modified gating properties in TRPM3 mutants causing intellectual disability and epilepsy. Elife 9 (2020). 10.7554/eLife.57190

5 Zhao, S., Yudin, Y. & Rohacs, T. Disease-associated mutations in the human TRPM3 render the channel overactive via two distinct mechanisms. Elife 9 (2020). 10.7554/eLife.55634

6 Dyment, D. A. et al. De novo substitutions of TRPM3 cause intellectual disability and epilepsy. Eur J Hum Genet 27, 1611–1618 (2019). 10.1038/s41431-019-0462-x

7 Vandewauw, I. et al. A TRP channel trio mediates acute noxious heat sensing. Nature 555, 662–666 (2018). 10.1038/nature26137

8 Wagner, T. F. et al. Transient receptor potential M3 channels are ionotropic steroid receptors in pancreatic beta cells. Nat Cell Biol 10, 1421–1430 (2008). 10.1038/ncb1801

9 Thiel, G., Rubil, S., Lesch, A., Guethlein, L. A. & Rossler, O. G. Transient receptor potential TRPM3 channels: Pharmacology, signaling, and biological functions. Pharmacol Res 124, 92–99 (2017). 10.1016/j.phrs.2017.07.014

10 Vriens, J. et al. TRPM3 is a nociceptor channel involved in the detection of noxious heat. Neuron 70, 482–494 (2011). 10.1016/j.neuron.2011.02.051

11 Su, S., Yudin, Y., Kim, N., Tao, Y. X. & Rohacs, T. TRPM3 Channels Play Roles in Heat Hypersensitivity and Spontaneous Pain after Nerve Injury. J Neurosci 41, 2457–2474 (2021). 10.1523/JNEUROSCI.1551-20.2020

12 Held, K. et al. Activation of TRPM3 by a potent synthetic ligand reveals a role in peptide release. Proc Natl Acad Sci U S A 112, E1363–1372 (2015). 10.1073/pnas.1419845112

13 Krugel, U., Straub, I., Beckmann, H. & Schaefer, M. Primidone inhibits TRPM3 and attenuates thermal nociception in vivo. Pain 158, 856–867 (2017). 10.1097/j.pain.0000000000000846

14 Becker, L. L. et al. Primidone improves symptoms in TRPM3-linked developmental and epileptic encephalopathy with spike-and-wave activation in sleep. Epilepsia 64, e61–e68 (2023). 10.1111/epi.17586

15 Vriens, J. & Voets, T. Sensing the heat with TRPM3. Pflugers Arch 470, 799–807 (2018). 10.1007/s00424-017-2100-1

16 Clapham, D. E. & Miller, C. A thermodynamic framework for understanding temperature sensing by transient receptor potential (TRP) channels. Proc Natl Acad Sci U S A 108, 19492–19497 (2011). 10.1073/pnas.1117485108

17 Yeh, F., Jara-Oseguera, A. & Aldrich, R. W. Implications of a temperature-dependent heat capacity for temperature-gated ion channels. Proc Natl Acad Sci U S A 120, e2301528120 (2023). 10.1073/pnas.2301528120

18 Bandell, M., Macpherson, L. J. & Patapoutian, A. From chills to chilis: mechanisms for thermosensation and chemesthesis via thermoTRPs. Curr Opin Neurobiol 17, 490–497 (2007). 10.1016/j.conb.2007.07.014

19 Caterina, M. J., Rosen, T. A., Tominaga, M., Brake, A. J. & Julius, D. A capsaicin-receptor homologue with a high threshold for noxious heat. Nature 398, 436–441 (1999). 10.1038/18906

20 Dhaka, A., Viswanath, V. & Patapoutian, A. Trp ion channels and temperature sensation. Annu Rev Neurosci 29, 135–161 (2006). 10.1146/annurev.neuro.29.051605.112958

21 Hensel, H. Thermoreception and temperature regulation. Monogr Physiol Soc 38, 1–321 (1981).

22 Talavera, K. et al. Heat activation of TRPM5 underlies thermal sensitivity of sweet taste. Nature 438, 1022–1025 (2005). 10.1038/nature04248

23 Patapoutian, A., Peier, A. M., Story, G. M. & Viswanath, V. ThermoTRP channels and beyond: mechanisms of temperature sensation. Nat Rev Neurosci 4, 529–539 (2003). 10.1038/nrn1141

24 Voets, T. et al. The principle of temperature-dependent gating in cold- and heat-sensitive TRP channels. Nature 430, 748–754 (2004). 10.1038/nature02732

25 Tan, C. H. & McNaughton, P. A. The TRPM2 ion channel is required for sensitivity to warmth. Nature 536, 460–463 (2016). 10.1038/nature19074

26 Song, K. et al. The TRPM2 channel is a hypothalamic heat sensor that limits fever and can drive hypothermia. Science 353, 1393–1398 (2016). 10.1126/science.aaf7537

27 McKemy, D. D., Neuhausser, W. M. & Julius, D. Identification of a cold receptor reveals a general role for TRP channels in thermosensation. Nature 416, 52–58 (2002). 10.1038/nature719

28 Peier, A. M. et al. A TRP channel that senses cold stimuli and menthol. Cell 108, 705–715 (2002). 10.1016/s0092-8674(02)00652-9

29 Story, G. M. et al. ANKTM1, a TRP-like channel expressed in nociceptive neurons, is activated by cold temperatures. Cell 112, 819–829 (2003). 10.1016/s0092-8674(03)00158-2

30 Caterina, M. J. Transient receptor potential ion channels as participants in thermosensation and thermoregulation. Am J Physiol Regul Integr Comp Physiol 292, R64–76 (2007). 10.1152/ajpregu.00446.2006

31 Talavera, K., Nilius, B. & Voets, T. Neuronal TRP channels: thermometers, pathfinders and life-savers. Trends Neurosci 31, 287–295 (2008). 10.1016/j.tins.2008.03.002

32 Grimm, C., Kraft, R., Sauerbruch, S., Schultz, G. & Harteneck, C. Molecular and functional characterization of the melastatin-related cation channel TRPM3. J Biol Chem 278, 21493–21501 (2003). 10.1074/jbc.M300945200

33 Oberwinkler, J. & Philipp, S. E. Trpm3. Handb Exp Pharmacol 222, 427–459 (2014). 10.1007/978-3-642-54215-2_17

34 Naylor, J. et al. Pregnenolone sulphate- and cholesterol-regulated TRPM3 channels coupled to vascular smooth muscle secretion and contraction. Circ Res 106, 1507–1515 (2010). 10.1161/CIRCRESAHA.110.219329

35 Mulier, M. et al. Upregulation of TRPM3 in nociceptors innervating inflamed tissue. Elife 9 (2020). 10.7554/eLife.61103

36 Krivoshein, G., Tolner, E. A., Maagdenberg, A. V. D. & Giniatullin, R. A. Migraine-relevant sex-dependent activation of mouse meningeal afferents by TRPM3 agonists. J Headache Pain 23, 4 (2022). 10.1186/s10194-021-01383-8

37 Aloi, V. D. et al. TRPM3 as a novel target to alleviate acute oxaliplatin-induced peripheral neuropathic pain. Pain 164, 2060–2069 (2023). 10.1097/j.pain.0000000000002906

38 de Sainte Agathe, J. M., et al. Confirmation and Expansion of the Phenotype Associated with the Recurrent p.Val837Met Variant in TRPM3. Eur J Med Genet 63, 103942 (2020). 10.1016/j.ejmg.2020.103942

39 Gauthier, L. W. et al. Description of a novel patient with the TRPM3 recurrent p.Val837Met variant. Eur J Med Genet 64, 104320 (2021). 10.1016/j.ejmg.2021.104320

40 Kang, Q. et al. A Chinese patient with developmental and epileptic encephalopathies (DEE) carrying a TRPM3 gene mutation: a paediatric case report. BMC Pediatr 21, 256 (2021). 10.1186/s12887-021-02719-8

41 Burglen, L. et al. Gain-of-function variants in the ion channel gene TRPM3 underlie a spectrum of neurodevelopmental disorders. Elife 12 (2023). 10.7554/eLife.81032

42 Huang, Y., Fliegert, R., Guse, A. H., Lu, W. & Du, J. A structural overview of the ion channels of the TRPM family. Cell Calcium 85, 102111 (2020). 10.1016/j.ceca.2019.102111

43 Zhao, C. & MacKinnon, R. Structural and functional analyses of a GPCR-inhibited ion channel TRPM3. Neuron 111, 81–91 e87 (2023). 10.1016/j.neuron.2022.10.002

44 Ruan, Z. et al. Structures of the TRPM5 channel elucidate mechanisms of activation and inhibition. Nat Struct Mol Biol 28, 604–613 (2021). 10.1038/s41594-021-00607-4

45 Hu, J. et al. Physiological temperature drives TRPM4 ligand recognition and gating. Nature 630, 509–515 (2024). 10.1038/s41586-024-07436-7

46 Reeves, K. C., Shah, N., Munoz, B. & Atwood, B. K. Opioid Receptor-Mediated Regulation of Neurotransmission in the Brain. Front Mol Neurosci 15, 919773 (2022). 10.3389/fnmol.2022.919773

47 Patapoutian, A., Tate, S. & Woolf, C. J. Transient receptor potential channels: targeting pain at the source. Nat Rev Drug Discov 8, 55–68 (2009). 10.1038/nrd2757

48 Kwon, D. H. et al. Heat-dependent opening of TRPV1 in the presence of capsaicin. Nat Struct Mol Biol 28, 554–563 (2021). 10.1038/s41594-021-00616-3

49 Nadezhdin, K. D. et al. Structural mechanism of heat-induced opening of a temperature-sensitive TRP channel. Nat Struct Mol Biol 28, 564–572 (2021). 10.1038/s41594-021-00615-4

50 Yin, Y. et al. Molecular basis of neurosteroid and anticonvulsant regulation of TRPM3. Nat Struct Mol Biol (2025). 10.1038/s41594-024-01463-8

51 Goehring, A. et al. Screening and large-scale expression of membrane proteins in mammalian cells for structural studies. Nat Protoc 9, 2574–2585 (2014). 10.1038/nprot.2014.173

52 Zheng, S. Q. et al. MotionCor2: anisotropic correction of beam-induced motion for improved cryo-electron microscopy. Nat Methods 14, 331–332 (2017). 10.1038/nmeth.4193

53 Rohou, A. & Grigorieff, N. CTFFIND4: Fast and accurate defocus estimation from electron micrographs. J Struct Biol 192, 216–221 (2015). 10.1016/j.jsb.2015.08.008

54 Scheres, S. H. RELION: implementation of a Bayesian approach to cryo-EM structure determination. J Struct Biol 180, 519–530 (2012). 10.1016/j.jsb.2012.09.006

55 Bepler, T., Kelley, K., Noble, A. J. & Berger, B. Topaz-Denoise: general deep denoising models for cryoEM and cryoET. Nat Commun 11, 5208 (2020). 10.1038/s41467-020-18952-1

56 Punjani, A., Rubinstein, J. L., Fleet, D. J. & Brubaker, M. A. cryoSPARC: algorithms for rapid unsupervised cryo-EM structure determination. Nat Methods 14, 290–296 (2017). 10.1038/nmeth.4169

57 Emsley, P., Lohkamp, B., Scott, W. G. & Cowtan, K. Features and development of Coot. Acta Crystallogr D Biol Crystallogr 66, 486–501 (2010). 10.1107/S0907444910007493

58 Afonine, P. V. et al. Real-space refinement in PHENIX for cryo-EM and crystallography. Acta Crystallogr D Struct Biol 74, 531–544 (2018). 10.1107/S2059798318006551

59 Chen, V. B. et al. MolProbity: all-atom structure validation for macromolecular crystallography. Acta Crystallogr D Biol Crystallogr 66, 12–21 (2010). 10.1107/S0907444909042073

60 Pettersen, E. F. et al. UCSF ChimeraX: Structure visualization for researchers, educators, and developers. Protein Sci 30, 70–82 (2021). 10.1002/pro.3943

